# CRISPR-Cas guided mutagenesis of chromosome and virulence plasmid in *Shigella flexneri* by cytosine base editing

**DOI:** 10.1101/2022.03.04.482438

**Authors:** Atin Sharma, Ruqiya Omer Aden, Andrea Puhar, David A. Cisneros

## Abstract

*Shigella* is a Gram-negative bacterium that invades the human gut epithelium. The resulting infection, shigellosis, is the deadliest bacterial diarrheal disease. Much of the information about the genes dictating the pathophysiology of *Shigella*, both on the chromosome and the virulence plasmid, was obtained by classical reverse genetics. However, technical limitations of the prevalent mutagenesis techniques restrict the generation of mutants in a single reaction to a small number, preventing large scale targeted mutagenesis of *Shigella* and the subsequent assessment of phenotype. We adopted a CRISPR-Cas dependent approach, where a nickase Cas9 and cytidine deaminase fusion is guided by sgRNA to introduce targeted C→T transitions, resulting in internal STOP codons and premature termination of translation. In proof-of-principle experiments using an *mCherry* fluorescent reporter, we were able to generate loss-of-function mutants in both *E. coli* and *Shigella* with up to 100% efficacy. Using a modified fluctuation assay, we determined that under optimized conditions, the frequency of untargeted mutations introduced by the Cas9-deaminase fusion is in the same range as spontaneous mutations, making our method a safe choice for bacterial mutagenesis. Further, we programmed the method to mutate well-characterized chromosomal and plasmid-borne *Shigella* genes and found the mutant phenotype to be similar to that of the reported gene deletion mutants, with no apparent polar effects at the phenotype level. This method can be used in a 96-well plate format to increase the throughput and generate an array of targeted loss-of-function mutants in a few days.

**Importance:** Loss-of-function mutagenesis is critical in understanding the physiological role of genes. Therefore, high-throughput techniques to generate such mutants are important for facilitating the assessment of gene function at a pace that matches system biology approaches. However, to our knowledge, no such method was available for generating an array of single gene mutants in an important enteropathogen - *Shigella*. This pathogen causes high morbidity and mortality in children and antibiotic resistant strains are quickly emerging. Therefore, determination of the function of unknown *Shigella* genes is of utmost importance to develop effective strategies to control infections. Our present work will bridge this gap by providing a rapid method for generating loss-of-function mutants. The highly effective and specific method has the potential to be programmed to generated multiple mutants in a single, massively parallel, reaction. By the virtue of plasmid compatibility, this method can be extended to other members of Enterobacteriaceae.

## Introduction

***S**higella* is a Gram-negative bacterium that belongs to the Enterobacteriaceae family. *Shigella flexneri* is the most prevalent species resulting in the disease – shigellosis or bacillary dysentery (1). Shigellosis is mostly a self-limiting disease in healthy adults but poses a major threat to children, elderly and the immunocompromised (2). With an annual death toll of 212,000, of which 63,713 are children below the age of 5, shigellosis is the deadliest bacterial diarrheal disease (3). Very low infection doses (10-100 bacteria), easy person-to-person spread and the acute inflammatory response are responsible for such high rates of incidence and mortality, especially in low- and medium-income countries (4). Being an invasive enteric pathogen, *Shigella* gains access to the intestinal epithelial cells by attaching and inducing its own uptake. Once inside the cells, it lyses the vacuole, replicates, and by using actin-based motility, disseminates to neighboring cells (reviewed in (5)). Virulence of *Shigella* requires a virulence plasmid-encoded type III secretion system (T3SS) which injects proteins into the host cell aiding the bacterium in establishment of infection through invasion, subversion of host defenses, and dissemination (reviewed in (6)).

**W**e recently resequenced the genome of *S. flexneri* serotype 5a M90T, a widely used lab reference strain, and mapped global transcription start sites (7). Unsurprisingly, many transcription start sites led to identification of genes that await functional characterization to reveal new and interesting information about *Shigella* physiology. Loss-of-function mutagenesis, and subsequent assessment of phenotype, is a common and perhaps the most convenient way to determine the function of unknown genes. The classic allele replacement technique, first described decades ago by Datsenko *et al*. (8), is still the method of choice for targeted mutations in *Shigella* (9–11), but it has some limitations. Since the success of this technique is directly dependent on transformation of a linear DNA construct, the requirement of a large amount of DNA and electroporation limit its throughput. The two-step process, allele replacement and subsequent removal, leaves a scar sequence which, if not optimized for being in-frame, results in polar effects that affect phenotype assessment (12). Although PCR-mediated construction of a linear DNA construct is easy, it requires expensive long primers and electroporation which adds to the final cost and acts as a technical bottleneck to increasing the throughput of mutagenesis.

**C**RISPR-Cas, a bacterial defense system against phages, has gained much attention for its use in genetic modifications (13). The *Streptococcus pyogenes* Cas9, the most commonly used protein for genome engineering, can be directed using guide RNA (gRNA) to introduce double-stranded (ds) DNA breaks (14). In eukaryotes, this is subsequently followed by DNA repair, resulting in generation of mutants, but in prokaryotes, a dsDNA break is lethal (15). Therefore, CRISPR-Cas based mutagenesis became the method of choice for targeted mutations in the eukaryotes, but its usage in prokaryotes was initially restricted to increasing the overall efficiency of allele replacement by positive selection (16, 17). However, later modifications resulted in the advent of base editors or catalytically impaired Cas9 proteins fused to deaminases, resulting in base substitution and, eventually, loss-of-function mutations (18, 19). These methods avoid dsDNA breaks and result in markerless mutations. Moreover, the requirement of plasmid borne sgRNA and effector proteins (Cas9 variants) eliminates the need of electroporation, thereby increasing the efficacy of the process, making it possible to multiplex or generate massively parallel mutations (20–22).

**H**ere, we describe a CRISPR-Cas mediated base editing method, using an established cytidine base editor (20, 22) that can be programmed to generate loss-of-function mutations in *S. flexneri*. By mutating well-characterized *S. flexneri* genes of known phenotype/function, encoded on both the virulence plasmid and the chromosome, we show that the one-step method is easy, fast, and highly effective in inactivating genes. Utilizing this method, multiple genes in *S. flexneri* can be mutated in parallel, showing its great potential for high-throughput mutagenesis.

## Results

### Construction of a two plasmid-based system for expression of sgRNA and nCas9-AID

**A**s an effector protein, we selected a fusion of catalytically inactive Cas9 - nickase Cas9 (nCas9^D10A^) and an ortholog of activation-induced cytidine deaminase (AID), previously described as an effective base editor in the Gram-positive industrial bacterium *Corynebacterium glutamicum* (22). The fusion protein is guided to the target gene by sgRNA (single guide RNA – 20 nt target-specific ‘spacer’ fused to a gRNA scaffold) where, within a window of −20 to −16 from the protospacer adjacent motif (PAM), the AID deaminates cytosine (C) to Uracil (U) and the nCas9 nicks the unmodified strand, resulting in its replacement and C→T (G→A on opposite strand) transition (Fig. 1A). This leads to conversion of internal CAG, CGA, and CAA codons on the coding strand or CCA on the non-coding strand into TAG, TGA, and TAA (STOP codons), premature termination of translation, and loss-of-function mutation.

**Figure 1:**
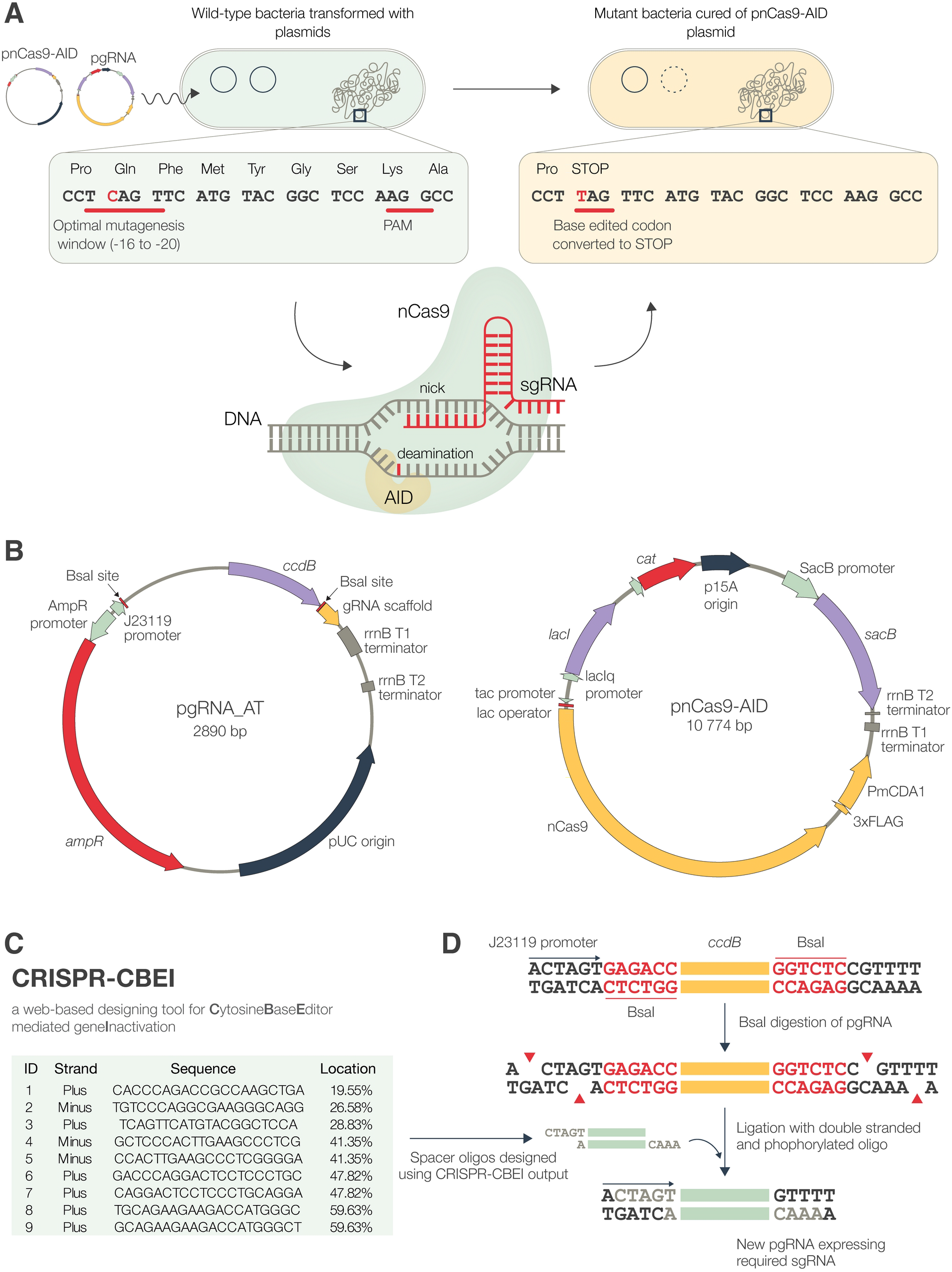
Overview of the CRISPR-Cas guided base editing method. (A) Generation of mutants using *mCherry* as example. After co-transformation with pnCas9-AID and pgRNA, expression of Cas9-AID and a specific sgRNA introduces a premature stop codon in the target sequence. The PAM and the editable window are underlined with the target C nucleotide shown in red. (B) Maps of plasmids pgRNA_AT and pnCas9-AID used for mutagenesis in *E. coli* and *S. flexneri*. (C) Output from CRISPR-CBEI for *mCherry* and (D) the general scheme for high-efficiency cloning of the spacers determined by CRISPR-CBEI.

**W**e generated a medium-copy plasmid based on pSU19 (23), pnCas9-AID, to express nCas9-AID (Fig. 1B). The SacB encoding gene was cloned to facilitate curing of this plasmid (24). To express the sgRNA, we constructed a high-copy plasmid, pgRNA_AT, that uses a synthetic J23119 promoter. The *tac* promoter, used to drive the expression of nCas9-AID, is already known to function in *Shigella* (25). However, to determine J23119 activity in *Shigella*, superfolder GFP (sfGFP) with a synthetic ribosome binding site was cloned in pgRNA_AT (pgRNA-(R)-sfGFP) and transformed into *S. flexneri* (Supplementary Fig. S1). Expression of sfGFP, visualized by green fluorescence, showed constitutive activity of the J23119 promoter in *S. flexneri*.

**T**o program a desired mutation, we used an online program - CRISPR-CBEI (26). It determines mutable sites and lists spacers required for the desired mutation in a target gene (Fig. 1C). We synthesized spacers as oligos with overhangs complementary to the ends of BsaI digested pgRNA_AT to ensure directional cloning (Fig. 1D). The oligos were hybridized and phosphorylated prior to ligation and transformed in *E. coli* DH5α. All the transformants obtained carried the required recombinant plasmid as the cells transformed with non-recombinant plasmid are killed (22, 27).

### Base editing effectively inactivates chromosomal genes in *E. coli* and *S. flexneri*

**W**ith optimal tools in hand, we tested our method by mutating a gene with an easy-to-read phenotype in *S. flexneri* and *E. coli* as a benchmark. We generated *attTn7::mCherry* insertion in *E. coli* MG1655 (*E. coli····mCherrý*) and *S. flexneri* M90T 5a (*S. flexneri::mCherry*) using Tn7 transposition. We generated pgRNA_m2, pgRNA_m3, and pgRNA_m4 to introduce premature STOP codons in mCherry at the 69^th^, 114^th^, and 98^th^ codon positions. We also constructed pgRNA-X, expressing a non-binding random spacer oligo.

**W**e co-transformed *E. coh::mC.herry* and *S. flexneri::mCherry* with pnCas9-AID or empty plasmid (pSU19) and with *mCherry*-targeting or control sgRNA plasmids (Fig. 2A). Co-transformants were selected as fluorescent colonies on selective medium containing glucose (Supplementary Fig. S2). To optimize the time and conditions necessary for mutagenesis, we plated actively growing cultures of *E. coh::mC.herry* and *S. flexneri::mCherry*, with or without IPTG induction. To determine the frequency of mutation (Fig. 2B, C), we counted the resulting colonies and scanned them for red fluorescence (Fig. 2D, E).

**Figure 2:**
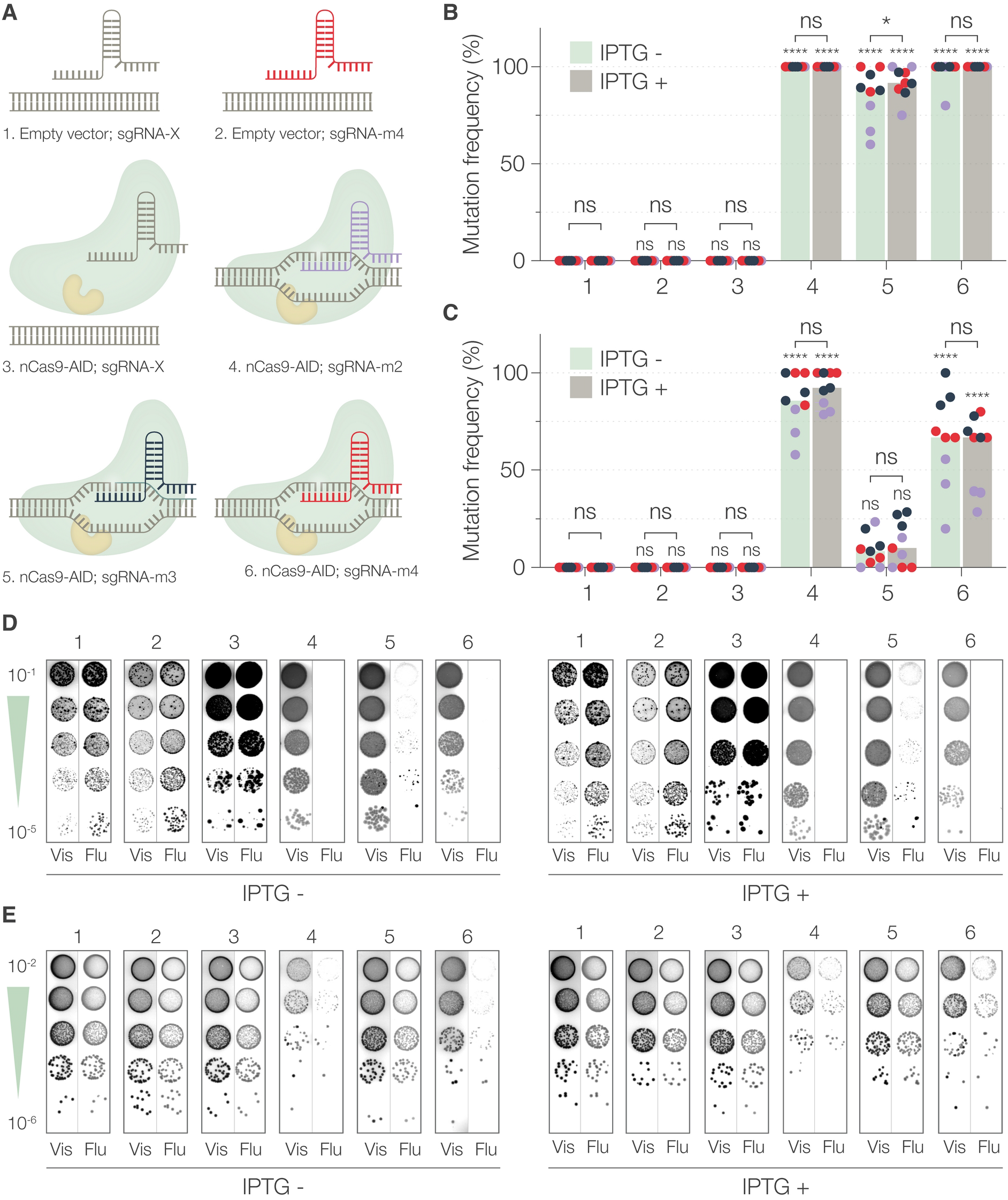
Mutagenesis of chromosomally encoded *mCherry* in *E. coli* and *S. flexneri*. (A) Schematic representation of the six different conditions (plasmid combinations), numbered from 1-6, used in the experiment. (B) Mutation frequency in *E. coli* and (C) *S. flexneri*, determined as the ratio of the number of non-fluorescent (mutant) colonies to the total number of colonies, after 2 hours of mutagenesis with or without IPTG induction. The numbers on the X-axis correspond to the conditions represented in (A). Each dot represents a technical replicate and is color-coded to represent a biological replicate. The bars represent the median of three independent experiments. Statistical significance was determined by performing two-way ANOVA followed by Sidák’s multiple comparison tests to determine the significance of each condition with respect to control conditions (condition 1) and to determine the significance of IPTG addition (ns – not significant, \**P*<0.0332, \*\**P*<0.0021, \*\*\**P*<0.0002, \*\*\*\**P*<0.0001). (D) Phenotypic determination of loss of fluorescence in *E. coli* and (E) *S. flexneri*. Representative visible (Vis) and fluorescence (Flu) images of colonies obtained by spotting logarithmic dilutions of induced (IPTG+) or not induced (IPTG-) cultures, after 2 hours. The numbers 1-6 correspond to the conditions represented in (A).

**I**n both species, the basal frequency of mutagenesis (no nCas9-AID and control gRNA-X) was zero (Fig. 2B-E, condition 1) and an *mCherry* targeting sgRNA alone (condition 2), or nCas9-AID with sgRNA-X, made no significant difference. This showed that gene-specific sgRNA alone or the nCas9-AID fusion bound to a control gRNA could not measurably introduce mutations. However, the frequency of mutagenesis was significantly higher than control, in both species, when nCas9-AID was expressed in presence of two out of three 5 *mCherry-targeting* sgRNA (Fig. 2B-E, conditions 4 and 6). Although the frequency was slightly lower in *S. flexneri* (median 85.7% and 66.7% as compared to 100% in *E. coli*), they were in the same order of magnitude irrespective of the addition of IPTG. This result shows that base-editing can be readily performed in *S. flexneri* with high efficiencies. However, a different effect was seen in case of sgRNA_m3. While the median frequency in *E. coli* was lower (87.8%) but in the same order as with sgRNA_m2 and sgRNA_m4, the median frequency in *S. flexneri* (8.3%) was not significantly different from the control condition (Fig 2C, condition 5). Addition of IPTG or longer incubation time did not improve the frequency (Fig. 2C, Supplementary Fig. S3). Together, these results indicated that sgRNAs had significant difference in their ability to target *mCherry* which shows species-dependent variation. Effective mutagenesis, however, could still be performed in both species, surprisingly, without chemical induction of nCas9-AID expression.

**W**e also determined the bacterial titer (CFU/ml) for all the time points in each condition (Supplementary Fig. S3). The overall titer increased with time in both species, but there was no significant correlation across timepoints compared to the control conditions (condition 1). Therefore, with the present data, we cannot strongly conclude if carrying pgRNA derivatives, with or without pnCas9-AID, have specific effects on the titers. Only a small, but measurable, effect could be attributed to the addition of IPTG, which results in changes in growth rates or viability.

### nCas9-AID is highly effective at undetectable levels

**D**ifferences in the efficacy of sgRNA_m3 in *E. coli* and *S. flexneri*, irrespective of IPTG induction, indicated that sgRNA, and not nCas9-AID expression, could be a factor responsible for the species-specific differences in the frequency of mutagenesis. To test this hypothesis, we transformed pgRNA-(R)-sfGFP into *E. coli* MG1655 and *S.flexneri::mScarlet (attTn7::mScarlet* in *S. flexneri*) and measured the sfGFP fluorescence over time. The fluorescence was higher in *E. coli* (up to 6-folds) as compared to fluorescence in *S. flexnerv::mScarlet* (Fig. 3A). This difference was confirmed at the single-cell level, when the green fluorescence was quantified by microscopy (Fig. 3B, Supplementary Fig. S4A). The expression of sfGFP from pgRNA-(R)-sfGFP was about four times higher in *E. coli* as compared to *S. flexneri* (Fig. 3C). This difference could be an outcome of several factors including, but not limited to, plasmid copy number, promoter activity, and sgRNA stability. This difference in sfGFP expression in the context of pgRNA_AT could explain differences in targeting efficiencies between *E. coli* and *S. flexneri*.

**Figure 3:**
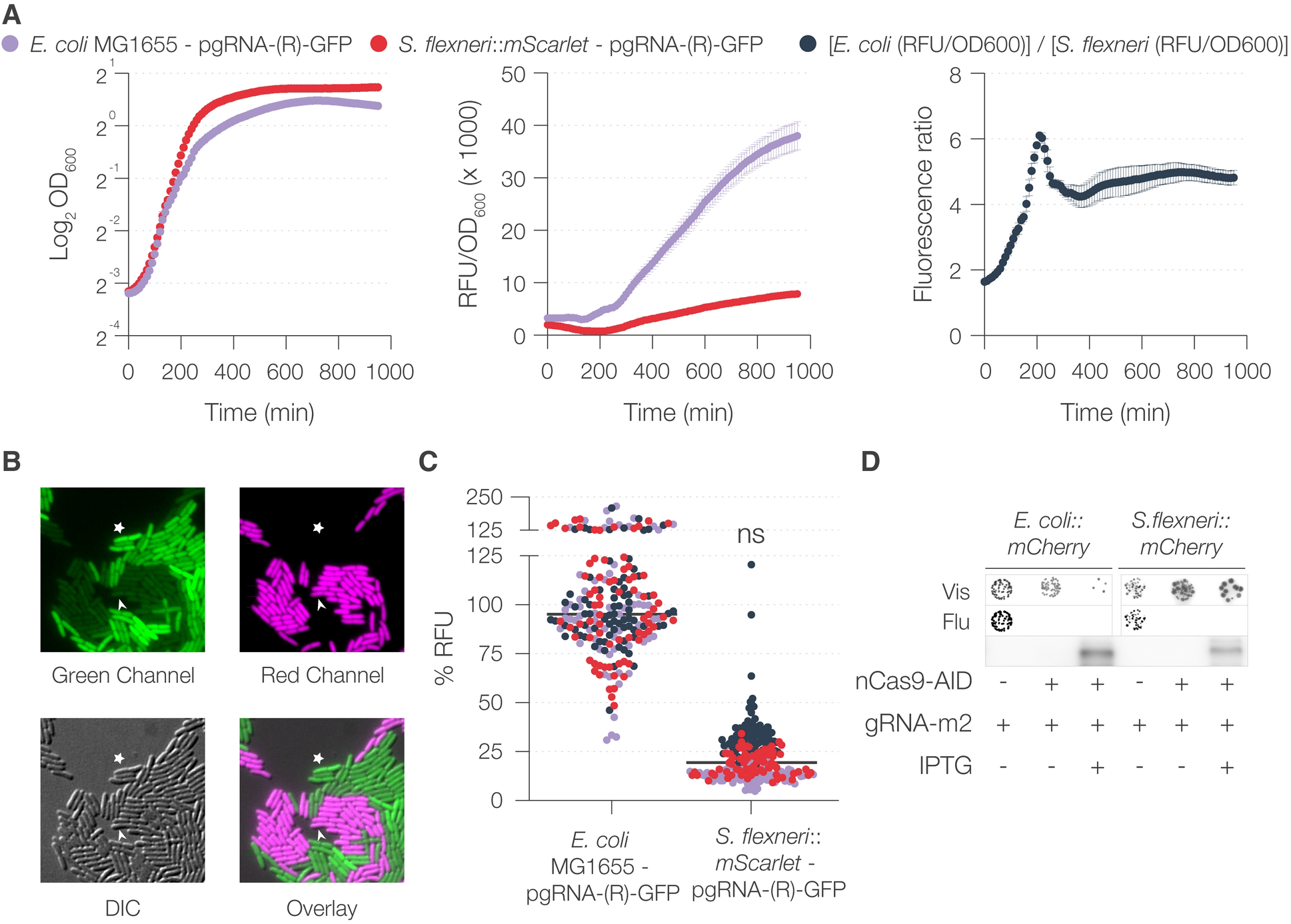
Comparison of expression from pgRNA and pnCas9-AID in *E. coli* and *S. flexneri*. (A) Comparison of sfGFP fluorescence from pgRNA-(R)-GFP in *E. coli* and *S. flexneri*::*mScarlet*. Fluorescence and OD600 of bacterial cultures were measured simultaneously in a multimode plate reader and the ratio (corrected fluorescence) was determined for each time point. Fold change in fluorescence was determined as a ratio of corrected fluorescence of *E. coli* and *S. flexneri::mScarlet* at each time point. (B) Representative image of a 50:50 mixture of the cultures as in (A) taken with a fluorescence microscope. The arrowhead denotes a representative *S. flexneri* bacterium and the asterisk marks a representative *E. coli* bacterium. (C) Quantification of fluorescence relative to the mean of *E. coli* fluorescence. The horizontal line represents the median. Each dot represents a technical replicate (fluorescent bacterium in one representative image) and is color coded to represent a biological replicate (image taken on different days). Statistical significance was determined by performing one-way ANOVA followed by Tukey’s multiple comparison tests (ns – not significant, \**P*<0.0332, \*\**P*<0.0021, \*\*\**P*<0.0002, \*\*\*\**P*<0.0001). (D) Expression of Cas9-AID fusion protein, determined by immunodetection using anti-Cas9 antibody. Mutagenesis of *mCherry*, guided by sgRNA_m2, was performed simultaneously. The outcome of the mutagenesis is shown as visible (Vis) and fluorescence (Flu) images of colonies obtained by spotting 10^-5^ dilution of the cultures.

**W**e had already determined that induction of the expression of nCAs9-AID was, surprisingly, not necessary for efficient mutagenesis. We wanted to determine if different levels of nCas9-AID correlated with *mCherry* mutagenesis guided by sgRNA_m2. Our results showed that in both *E. coli* and *S. flexneri*, *mCherry* was mutated despite nCas9-AID levels being below the limit of detection as determined by Western Blot (Fig 3D). nCas9-AID was detected upon IPTG induction, but accompanied by the appearance of bands below 100 kDa recognized by anti-Cas9 antibody, suggesting partial nCas9-AID degradation (Supplementary Fig. S4B). Interestingly, a previous study used dCas9-AID fused to a degron to decrease its expression (21). We seem to have achieved the same outcome, albeit inadvertently. These results indicate that the nCas9-AID fusion protein is not very stable but highly effective, nonetheless, even at barely detectable levels.

### nCas9-AID expressing strains are not hypermutators

**W**e suspected that leaky expression of nCas9-AID could alter the frequency of mutation at loci other than those targeted due to unspecific activity of the fused AID domain (28, 29). To test the basal mutation rate in *S. flexneri*, we performed the classic fluctuation assay using nalidixic acid (30). Point mutations in gyrase encoding gene, *gyrA*, render *Shigella* resistant to nalidixic acid (31). The frequency of appearance of such mutants can be used as a measure 7 of basal mutation rate. We designed a sgRNA (sgRNA-gyrA) that would cause a G→A transition at position 259, resulting in Asp_87_ → Asn_87_ substitution and resistance to nalidixic acid. *S. flexneri* strains carrying backbone plasmid pSU19, instead of pnCas9-AID, were used to distinguish between the mutation rate at the *gyrA* locus in presence or absence of nCas9-AID (Fig 4A). Mutagenesis was carried out as in case of *mCherry*, but in addition to spotting on TSA plates, 10-fold serial dilutions of the cultures were also spotted on TSA plates containing nalidixic acid and the resulting colonies were counted.

**Figure 4:**
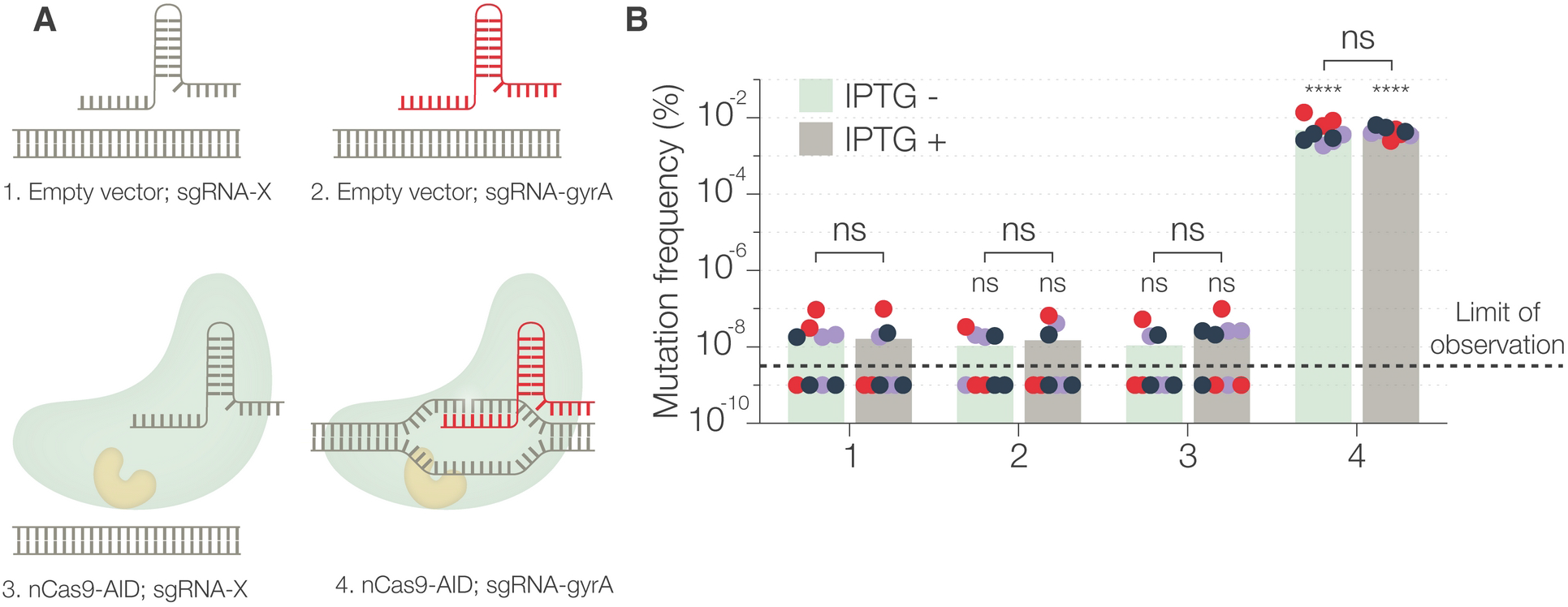
Nalidixic acid resistance assay to compare the frequency of guided and random mutations. (A) Schematic representation of the four different conditions (plasmid combinations), numbered 1-4, used in the experiment. (B) Mutation frequency, determined as the ratio of the number of nalidixic acid resistant (mutant) colonies to the total number of colonies, after 2 hours of mutagenesis with and without IPTG induction. The numbers on the X-axis correspond to the conditions represented in (A). Each dot represents a technical replicate and is color-coded to represent a biological replicate. In some technical replicates, the mutation frequency was below the limit of observation (1×10^-9^) as spotting high concentrations of the bacterial culture resulted in a bacterial mass, instead of distinct antibiotic resistant colonies. The bars represent the median of three independent experiments. Statistical significance was determined by performing two-way ANOVA followed by Sidák’s multiple comparison tests for each condition with respect to the control (condition 1) and for IPTG addition (ns – not significant, \**P*<0.0332, \*\**P*<0.0021, \*\*\**P*<0.0002, \*\*\*\**P*<0.0001).

**T**he background rate of *gyrA* mutation in absence of nCas9-AID, as a proxy measure of overall random mutation frequency, was in the order of 10^-8^ after 2 hours of incubation (Fig. 4B). It was not significantly altered when IPTG was added. Importantly, the expression of nCas9-AID, in presence of a non-targeting sgRNA (sgRNA-X), also did not significantly affect the mutation frequency. Even at later time points, the frequency of mutation for strains expressing nCas9-AID with sgRNA-X was similar to those not expressing nCas9-AID at all (Supplementary Fig. S4C). It was only in the presence of a *gyrA*-specific sgRNA (sgRNA-gyrA) that nCas9-AID was able to introduce nalidixic acid resistance mutations at a higher frequency. A nearly 5 log-fold increase in the frequency of mutation confirmed the specificity of our method. The frequency of mutation, however, was not as high as with *mCherry* mutagenesis, highlighting again that efficacy can depend on target genes and sgRNAs. The low-level leaky expression, and even induced expression of nCas9-AID, was not mutagenic by itself, probably due to the low stability of the fusion protein as observed by Western blot. This also suggested that the probability of introduction of unwanted unspecific mutations by nCas9-AID, under the conditions we tested, was in the same order of magnitude as spontaneous mutations.

### Loss-of-function mutants generated by base editing have the same phenotype as gene deletion mutants

**E**ncouraged by our results, we decided to use our system to generate mutations in parallel in well-characterized virulence genes on both the chromosome and the virulence plasmid and verify their phenotype (Table 1, Fig. 5A). We mutated *mxiD* in *S. flexneri* and performed a gentamicin protection assay to assess whether it had the same impact on invasion as reported earlier (32, 33). Wild-type and *S. flexneri mxiD(Q323X)* (Q_323_ residue substituted with STOP codon) were allowed to infect TC7 intestinal epithelial cells *in vitro* and the extracellular bacteria were killed by gentamicin treatment. Bacterial invasion was quantified by lysing the cells, to release intracellular bacteria that were protected from gentamicin, and spotting log-fold dilutions of the lysates on TSA plates (Fig. 5B). WT bacteria successfully invaded TC7 cells, but we did not observe any invasion by *S. flexneri mxiD(Q323X)* (Fig. 5C). Since *mxiD* mutants fail to assemble a functional T3SS, they are unable to secrete the T3SS substrates (32). Indeed, we did not recover any secreted proteins in the supernatant of *S. flexneri mxiD(Q323X)*, induced for secretion by Congo red (Fig 5D) (34).

**Figure 5:**
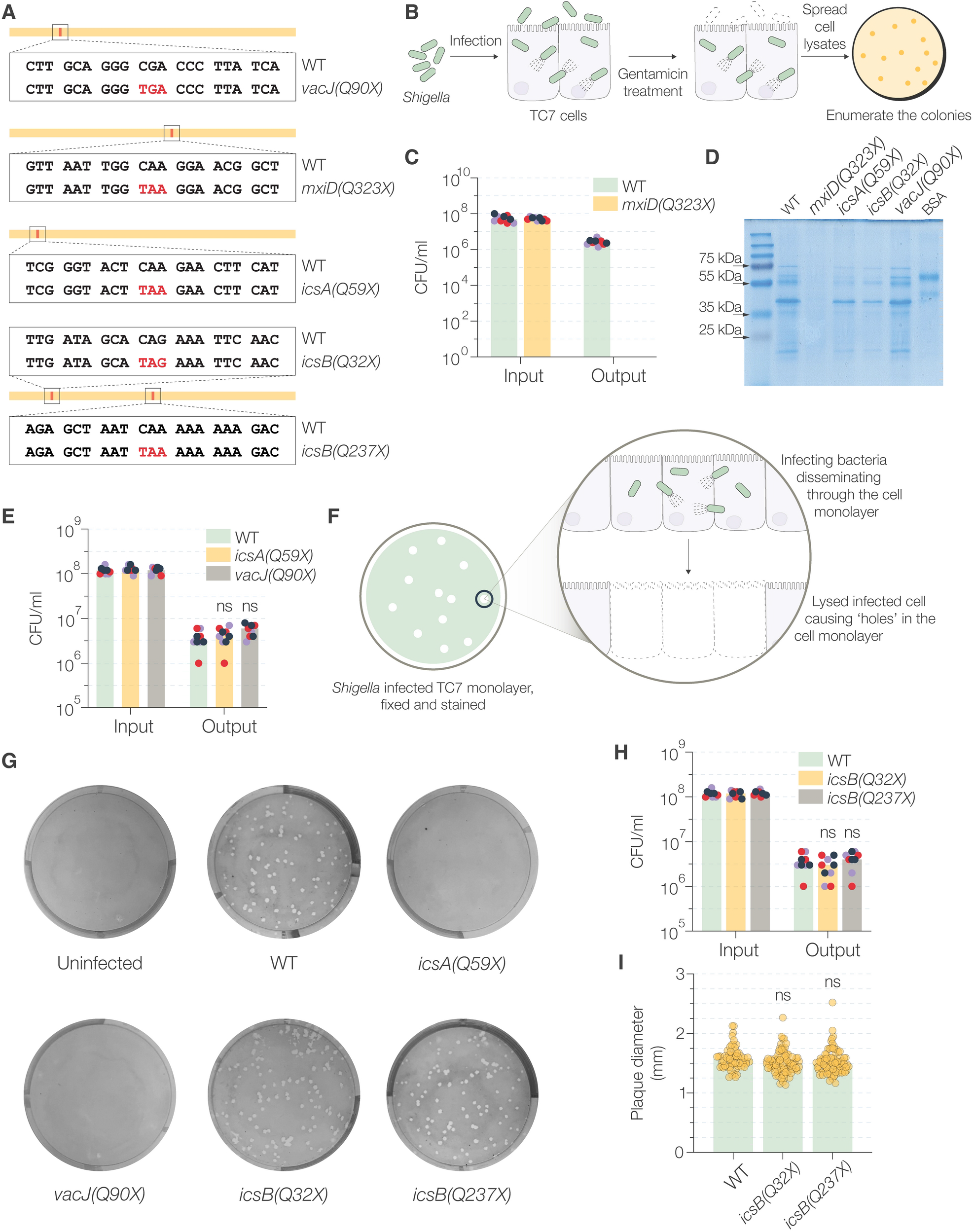
CRISPR/Cas-guided mutagenesis of characterized *S. flexneri* genes and assessment of their phenotype. (A) Representation of the STOP codons (highlighted in red) introduced by base-editing. The total length of the gene is indicated by a line and the red mark indicates the location of the mutable target site. (B) Schematic representation of the gentamicin protection assay used to assess the invasiveness of *Shigella*. (C) Invasiveness of *S. flexneri* WT and *mxiD(Q323X)*, determined by the number of bacterial CFU used to infect TC7 cells (input) and the number of bacterial CFU that were successful at invading them (output). In case of *S. flexneri mxiD(Q323X)*, the CFU count was not above the technical limit of observation (10^2^ CFU/ml). (D) SDS-PAGE gel showing the total secreted proteins precipitated from the supernatants of indicated bacterial cultures after induction of secretion by Congo red. BSA was used as a control for protein precipitation. (E) Invasiveness of WT, *icsA(Q59X)*, and *vacJ(Q90X) S. flexneri* determined by the gentamicin protection assay. (F) Schematic representation of the plaque formation assay. (G) Intracellular dissemination of WT and mutant *S. flexneri* determined by plaque formation assay. The images depict the stained monolayer of TC7 cells in a 6-well plate with the plaques seen as zones of clearance. (H) Invasiveness of WT and the two strains of *S. flexneri icsB* mutants determined by gentamicin protection assay. (I) Measurement of the diameters of the plaques from one representative plaque formation assay. In all the plots, each dot represents a technical replicate and is color-coded to represent a biological replicate. The bars represent the median of three independent experiments (except in (I)). Statistical significance was determined by performing two-way ANOVA followed by Sidák’s multiple comparison test (ns – not significant, \**P*<0.0332, \*\**P*<0.0021, \*\*\**P*<0.0002, \*\*\*\**P*<0.0001).

**Table 1.**
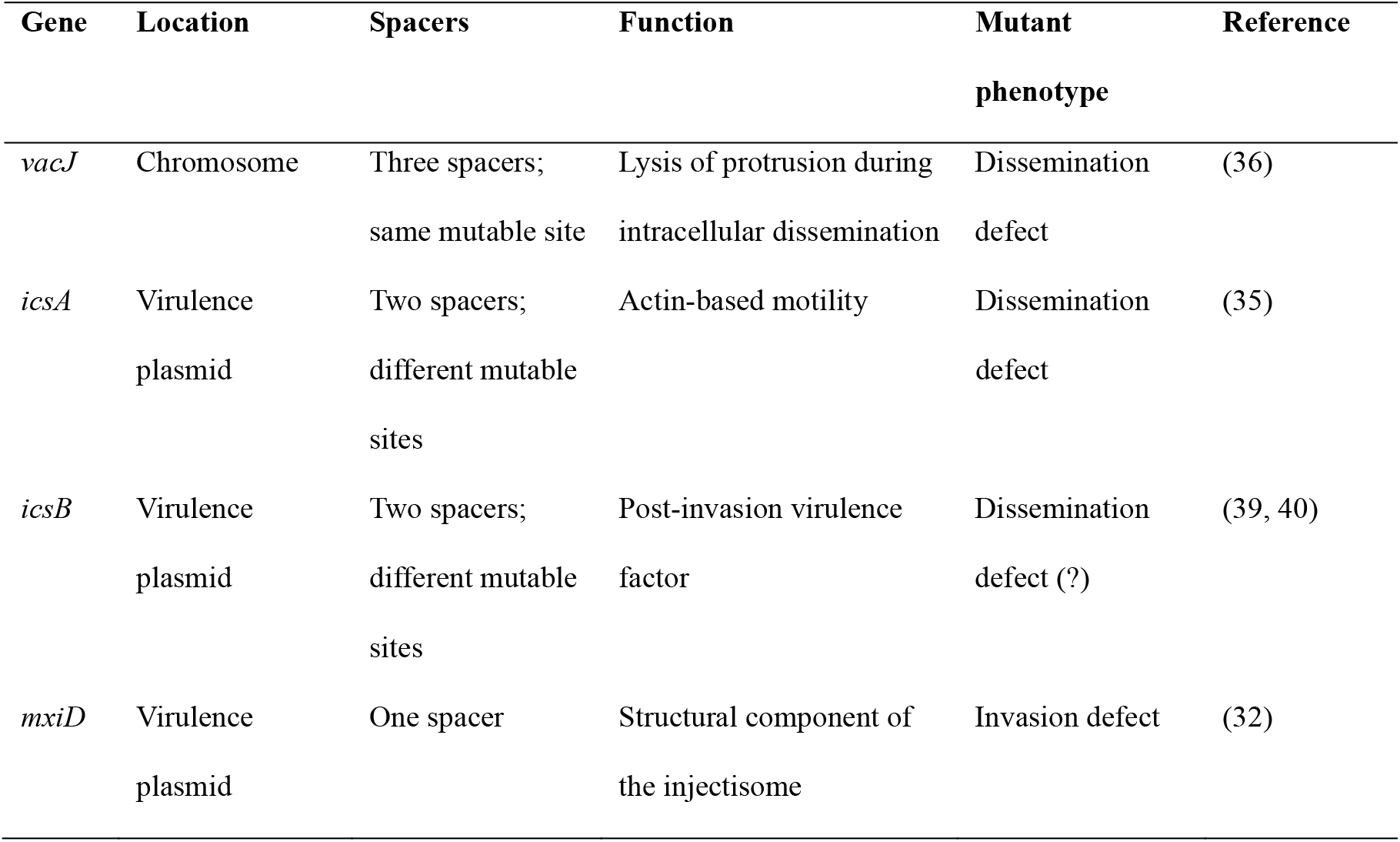
Details of characterized *S. flexneri* genes that were selected for mutagenesis. The selected genes are found both on the virulence plasmid and the chromosome. The variation in the number of spacers and their corresponding mutable sites is also mentioned.

***i**csA* and *vacJ* have been reported to be necessary for intracellular dissemination of *Shigella*, but are not involved in invasion (35, 36). We first verified that base-edited mutants *S. flexneri icsA(Q59X)* and *S. flexneri vacJ(Q90X)* had no invasion defect (Fig. 5E). To assess intracellular dissemination, we used the plaque formation assay, where cells are infected with bacteria but the bacteria can only disseminate intracellularly due to the presence of gentamicin in the medium (37, 38). Successful intracellular dissemination results in spread of bacteria and subsequent cell death, which appears as a zone of clearance in the cell monolayer (Fig 5F). Successful invasion and subsequent dissemination were evident in case of WT *S. flexneri* as numerous plaques of similar diameter were observed (Fig. 5G). However, no plaques were visible in case of *S. flexneri icsA(Q59X)* and *S. flexneri vacJ(Q90X)*, even at higher MOIs, indicating an obvious dissemination defect.

**T**aken together, these results showed that the phenotype of the mutants generated by our base editing method was consistent with that reported in the literature.

### Base editing results in phenotypically non-polar mutations

Polar effects, or decreased expression of genes downstream of the site of mutation, are a common mutagenesis problem in bacteria for polycistronic transcriptional units (12). For example, polar effects of *icsB* mutation have been shown to impair intracellular dissemination in *Shigella* by affecting transcription of downstream *ipa* genes (9, 39, 40). We generated two *icsB* mutants (*S. flexneri icsB(Q32X)* and *icsB(Q237X))* with the premature STOP codon introduced at two different sites (Fig. 5A). The *icsB* mutants had no apparent invasion defect (Fig. 5H) and there was no significant difference in the number and size of the plaques as compared to WT bacteria (Fig. 5G and 5I). This indicated that base edited *icsB* mutant might be non-polar.

To verify this effect at the mRNA level, we additionally mutated *ipgB*, a gene upstream *ipaC*, by constructing two *ipgB* mutant strains [*S. flexneri ipgB(Q44X)* and *S. flexneri ipgB(Q190X)}*. We determined the expression of *icsB, ipgB* and *ipaC* by qPCR (Fig. 6A). Two-way analysis ANOVA showed no significant differences between the transcription of *ipgB* and *ipaC* between wild-type *S. flexneri* and the four *icsB* and *ipgB* mutants (Fig. 6B). Moreover, no mutation showed significant differences compared to wild-type *S. flexneri* when transcription of *ipgB* or *ipaC* was analyzed individually (Supplementary Fig. S4D). As expected, no effect was observed at the protein level, assessed by comparing the secretion of IpaC (Fig. 6C). Although a loss-of-function mutation of *ipgB* has been reported to be 50% less invasive (41), we did not observe this decrease, mainly because this reduction is within experimental error in our hands (Fig. 6D). We did observe a significant effect on the levels of *icsB* transcription (Supplementary Fig. S4D), particularly for the *S. flexneri icsB(Q32X)* mutant, which is consistent with our previous report (42) that some mutations, including premature STOP codons, can decrease mRNA levels without affecting observed phenotypes. Altogether, these results showed that base-edited premature STOP codon mutations did not result in any observable polar effects, at least at the phenotype level.

**Figure 6:**
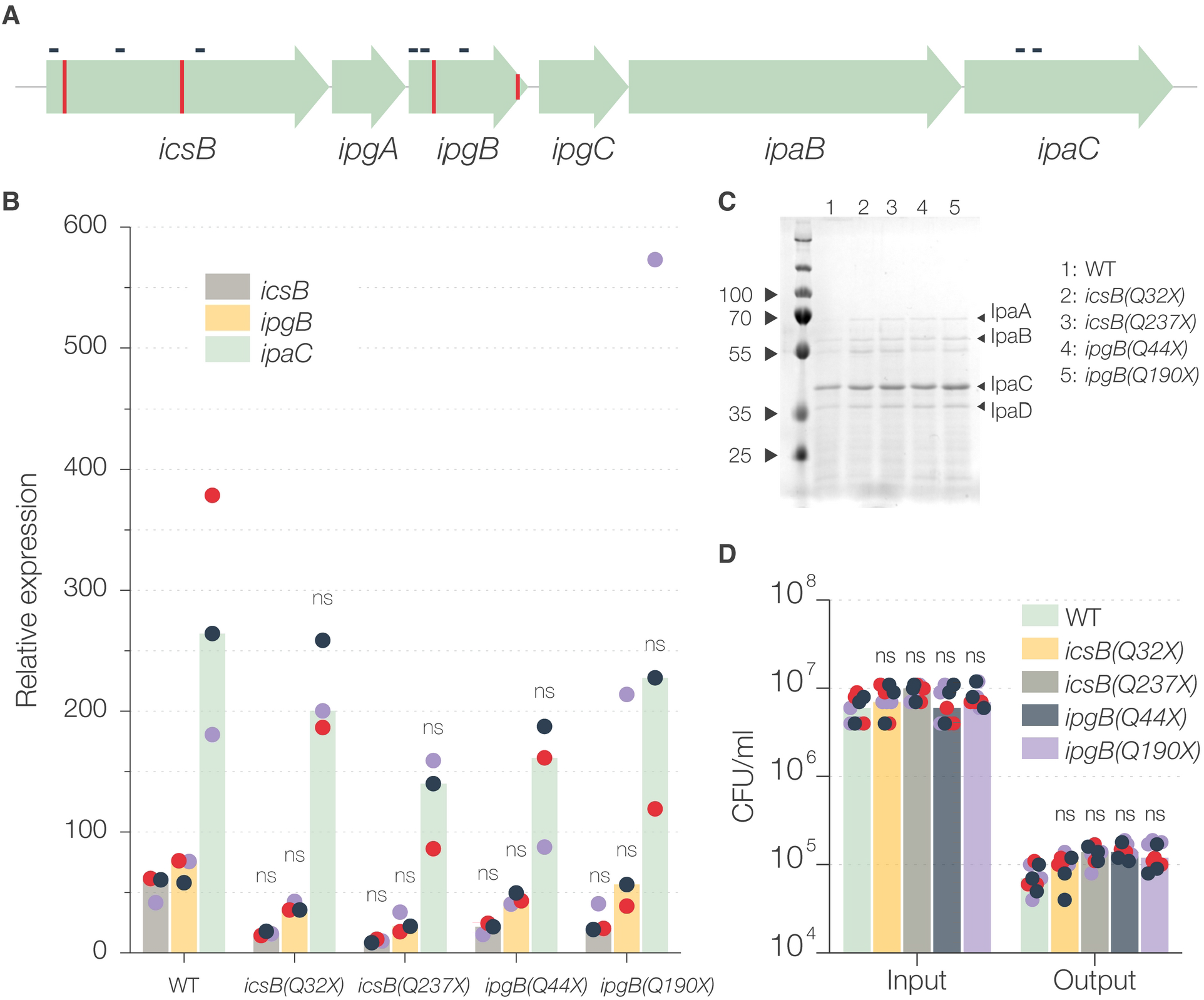
Assessment of polar effects of base-edited mutations. (A) Representation of the *icsB* operon. The red vertical lines represent the site of introduction of STOP codon in *icsB* and *ipgB* genes. The small horizontal black lines represent the amplicon used to determine the mRNA abundance by qPCR. (B) Normalized (to *cysG* and *hcaT*) expression of *icsB*, *ipgB*, and *ipaC* genes in different mutant strains. Each dot represents a single amplicon and is color-coded to represent a biological replicate. Statistical significance was determined by performing a two-way ANOVA and Holm-Sidák’s multiple comparison tests compared to WT (ns – not significant, \**P*<0.0332, \*\**P*<0.0021, \*\*\**P*<0.0002, \*\*\*\**P*<0.0001). (C) SDS-PAGE gel showing the secreted proteins (labelled with arrowheads) precipitated from the supernatants of indicated bacterial cultures after induction of secretion by Congo red. (D) Invasiveness of *S. flexneri* strains, determined by the number of bacterial CFU used to infect TC7 cells (input) and the number of bacterial CFU that were successfully invaded (output). Each dot represents a technical replicate and is color-coded to represent a biological replicate. The bars represent the median of three biological replicates. Statistical significance was determined by performing a two-way ANOVA followed by Sidák’s multiple comparison test compared to WT (ns – not significant, \**P*<0.0332, \*\**P*<0.0021, \*\*\**P*<0.0002, \*\*\*\**P*<0.0001).

## Discussion

**F**unctional characterization of genes is a primary step in understanding the physiology and pathogenesis of microbes, which is made possible by powerful tools such as high-throughput reverse genetics (43). However, the success of this approach is entirely dependent on the ability to generate loss-of-function libraries with an equally high throughput (44). Using a two-plasmid system and substituting electroporation with chemical transformation we were able to increase throughput ensuring pure colonies of several desired mutants prepared in parallel, in just 5-6 days from the receipt of spacer oligos (Supplementary Fig. S5). The use of type II restriction enzyme BsaI not only ensures directional cloning of spacer oligos but also provides the advantage that the parent plasmid needs to be digested only once for all cloning reactions. Furthermore, even for the gRNA that showed no significant frequency of mutation (sgRNA_m3) compared to the control, the reaction yielded a workable number of mutant colonies.

**A**lthough high frequency of mutation was achieved in previous studies by combining the expression of catalytically inactive Cas9, dCas9-AID, and a uracil DNA glycosylase inhibitor (21, 46), we avoided using this combination as the uracil DNA glycosylase inhibitor can potentially induce hypermutation, owing to its ability to impair DNA repair (47). In our case, the apparent low stability of nCas9-AID and combined expression with a sgRNA, targeting or non-targeting, might have decreased the availability of free apo-nCas9-AID, preventing it from binding to non-specific sites on genomic DNA (48) and introducing unwanted, potentially toxic mutations. Since the virulence plasmid is not necessary for bacterial survival, improved yield of mutants by counter-selecting the non-modified cells using CRISPR-Cas9 and recombineering methods together, is not applicable to mutation of plasmid-borne genes (16, 17). However, we were able to mutate plasmid and chromosomal genes in *S. flexneri* in parallel using the same strategy and workflow.

**I**t is now clear that polar effects are prevalent even in well-characterized systems such as the Keio collection (49). Therefore, even if we did not observe any polar effects at the phenotype level in this study, we observed potential changes in mRNA abundance and recommend that such effects are taken into account while generating base-edited mutants, in equal manner as for any other mutagenesis strategy.

**I**n addition to generating parallel single gene mutations, a spacer oligo library can be cloned into pgRNA in a single reaction resulting in a pgRNA plasmid library. This library can be transformed into bacteria expressing nCas9-AID, generating an equally large pooled library of loss-of-function mutants. Since the pgRNA (ColE1-derived origin) propagates stably (50), cloned spacer oligos can function as barcodes, aiding in high-throughput screening of gene function (51) and base editing can be subsequently used for targeted mutagenesis of selected candidates.

**D**espite many apparent advantages, the most obvious limitation of the base editing method is the availability of mutable sites in a particular gene. Certain genes, for example *mxiD* in this study, have only one mutable site. As seen from our results, certain sgRNAs are more effective than others and not having any alternative, therefore, can be limiting. However, with the discovery of Cas9 variants with varying PAM specificity, there is a possibility of combining them with deaminases to expand the arsenal of effector proteins to overcome this limitation (52). Altered PAM specificity can also address another possible limitation in form of the usage of amber codon for termination. Amber codon is only rarely used for termination of essential genes and is often a suboptimal STOP codon (53).

**I**n conclusion, we have successfully adapted a programmable CRISPR-Cas9 base editing method to generate loss-of-function mutants in *S. flexneri* and *E. coli*. This method can allow functional characterization of unknown *S. flexneri* genes by high-throughput reverse genetics and could be extended to other members of Enterobacteriaceae owing to plasmid and promoter compatibility.

## Methods

### Bacterial strains, cells, and culture conditions

**B**acterial strains, plasmids, and cells used in this study have been summarized in the Supplementary Table S1. All *E. coli* strains were grown in LB, *S. flexneri* in TSB, and TC7 cells were gown in DMEM containing 10% FBS, non-essential amino acids and penicillin-streptomycin. For maintenance of plasmids in *E. coli*, chloramphenicol and carbenicillin were used at 25 μg/ml and 100 μg/ml, respectively. In case of *S. flexneri*, the concentration was 15 μg/ml and 50 μg/ml, respectively. Spectinomycin was used at 50 μg/ml. *S. flexneri* colonies were selected on agar plates supplemented with 0.01% Congo red.

### Construction of fluorescent strains

**F**luorescent strains of *E. coli* and *S. flexneri*, constitutively expressing *mCherry* (and *mScarlet-i* in case of *Shigella*) from the chromosome, were constructed using Tn7-mediated transposition (54, 55). The detailed methodology is mentioned in Supplementary Methods S1.

### Plasmid construction

**A**ll the oligos used for recombinant procedures are mentioned in Supplementary Table S2. pgRNA_AT was generated by combining parts from pgRNA-ccdB and a pBluescript SK variant that lacks the BsaI site in the β-lactamase gene (22). pnCas9-AID was generated by combining parts of pSU19 and pdCas9-AID by *in-vivo* assembly (IVA) cloning (22, 56). Plasmid construction is explained in detail in Supplementary Methods S1.

### Guide RNA design and cloning

**T**he guide RNA for mutagenesis were designed using the program CRISPR_CBEI (26) (Supplementary Table S2). Details of cloning and guide RNA design are mentioned in Supplementary Methods S1.

### Mutagenesis

**p**nCas9-AID and respective pgRNA were co-transformed in ultra-competent *E. coli* using the heat shock method (57). The co-transformants were selected on LA plates supplemented with chloramphenicol, carbenicillin, and glucose. Since co-transformation was not very efficient in *S. flexneri*, pnCas9-AID and various sgRNA plasmids were sequentially transformed.

**B**acteria carrying both plasmids were grown overnight at 37°C in broth supplemented with chloramphenicol, carbenicillin, and glucose. The overnight culture was diluted 100 times in fresh medium and the bacteria were grown till OD_600_ = 0.2. 1 mM IPTG was added to overexpress nCas9-AID and the cultures were incubated at 37°C for further 2 hours (longer incubation times were tested in case of *mCherry* and *gyrA* mutagenesis). 10-fold serial dilutions of the cultures were made in sterile PBS and 10 μl of each dilution was spotted on agar plates containing no antibiotics. For *gyrA* mutagenesis, the agar plates were supplemented with nalidixic acid (30 μg/ml) and in addition to 10-fold dilutions, a 10-fold concentrated culture was also spotted. The agar plates were incubated overnight at 37°C and the colonies were enumerated. The plates were imaged in a Typhoon scanner (GE Healthcare) to determine fluorescence. Four random colonies were selected in each case to verify mutation by PCR and sequencing.

### Fluorescence measurement and microscopy

Overnight cultures of *E. coli* MG1655 and *S. flexneri::mScarlet* carrying pgRNA-(R)-GFP were diluted 100 times in fresh medium containing carbenicillin. 100 μl of culture was pipetted into different wells of a 96-well plate and the fluorescence (Ex: 480 nm and Em: 510 nm) and OD_600_ was measured simultaneously every 10 min in a SPARK multimode plate reader (Tecan) at 37°C. After 3 hours of incubation, 10 μl of both cultures were mixed in a tube and spotted onto a thin agarose pad on a glass slide. Microscopy images were taken using widefield microscope (Nikon) with DIC, Texas red, and FITC filters. The images were given false colors and fluorescence quantification was performed using ImageJ.

### Immunodetection of nCas9-AID

Bacteria were cultured similarly to the mutagenesis assay but instead of spotting dilutions on agar plates, 10 ml of cultures were pelleted by centrifugation and resuspended in 250 μl of 1X Laemmli buffer. The samples were boiled and resolved on 10% SDS-PAGE. The gels were stained using SimplyBlue stain (ThermoFisher) or used for transfer of proteins onto PVDF membranes (GE Healthcare). Probing was done using anti-Cas9 primary antibody (ThemoFisher, PA5-90171) at 1:20,000 dilution and HRP-conjugated anti-rabbit secondary antibody (Abcam) at 1:40,000 dilution. The blot was developed using ECL reagent (Cytiva) and imaged using Amersham imager 680 (Sigma Aldrich).

### Bacterial invasion assay and plaque formation assay

**T**he ability of bacteria to invade epithelial cells was determined by gentamicin protection assay as described earlier (33). All mutant *Shigella* strains were transformed with pAfaE, a derivative of pIL22 (58) expressing AFA I adhesin. Plaque formation assay was performed as described in (38) with modifications as follows. TC7 cells were infected at an MOI of 1:500 (bacteria:cells) and additionally at 1:150 for *vacJ* and *icsA* mutants. The infected cells were incubated for 48-72 h before fixation and staining.

### Secretion assay

**S**ecretion assays were performed as described earlier with modifications (59). Overnight cultures of bacteria were diluted 200 times and were grown till OD600 = 0.5. Cells were collected from 4 ml culture, by centrifugation, and washed twice with PBS. The pellet was finally resuspended in 1 ml prewarmed PBS containing 0.01% Congo red and incubated for 1 hour at 37°C. Bacteria were separated by centrifugation and the proteins in the supernatants were precipitated at 4°C for 10 min using 20% trichloroacetic acid. 1 mg/ml solution of BSA was used as a control for precipitation. The precipitated proteins were washed twice with ice-cold acetone and resuspended in Laemmli buffer before being resolved on 12% polyacrylamide gel. The gel was soaked overnight in 50% ethanol to remove Congo red and rehydrated gradually. The gel was stained with SimplyBlue SafeStain (ThermoFisher) to visualize the proteins.

### 96-well mutagenesis

**5**0 μl chemically competent *S. flexneri* carrying pnCas9-AID were pipetted into wells of a pre-chilled 96-well plate. sgRNA plasmids were added to individual wells and transformed by the heat-shock method. 150 μl of recovery medium (SOC) was added to each well and the plate was incubated at 37°C for 1 h. 100 μl of cell suspension and 1/10^th^ dilution from each well were spread onto selective plates. The plates were incubated overnight at 37°C and single colonies were picked up for mutagenesis. In a 96-well deep-well plate (Eppendorf), single colonies were inoculated in 500 μl of TSB with antibiotics and grown overnight at 37°C. 10 μl of overnight culture was added to 500 μl of fresh medium in a new 96-well deep well plate, which was incubated till OD_600_ = 0.2. 1 mM IPTG was added to the wells and the plate was incubated for another 2 h before serial dilutions (10 μl of 10^-5^ and 10^-6^ dilution) of the cultures were spotted on Congo red plates. The plates were incubated at 37°C overnight and single colonies were checked for desired mutation by colony PCR and DNA sequencing.

### qPCR analysis

**T**otal RNA extraction and qPCR were performed as previously described, with minor modifications (42, 60). Bacteria were grown similarly to the mutagenesis experiment but instead of spotting the dilutions, the cultures were treated with ice-cold Stop Solution (95% v/v ml of ethanol, 5% v/v ml of phenol pH 4) for 30 min on ice. The cells were then collected by centrifugation and stored at −80°C. cDNAs were subsequently prepared with random hexamer primers using a first strand cDNA synthesis kit (ThermoFisher). To analyze the 16 expression of *icsB, ipgB, ipaC*, and two housekeeping genes, *hcaT* and *cysG* (61), we designed specific oligonucleotides (Supplementary Table S2). To analyze qPCR data, we used the iQ5 optical system software (Bio-Rad) and obtained the Ct with default settings. We manually verified that changes in the threshold did not affect or bias the results. To normalize the data, the average Ct value of the two housekeeping genes was used to calculate a ΔCt for the gene of interest. Controls with no reverse transcriptase were also analyzed to exclude the presence of contaminating genomic DNA fragments.

## Acknowledgement

AP acknowledges funding from Swedish Research Council (Vetenskapsrådet) grant number #2016-06598, institutional support from Umeå University, Knut and Alice Wallenberg Foundation grant KAW 2015.0225, and Carl Kempe Foundation grant JCK-2031.3. DAC acknowledges support from the Carl Tryggers Stiftelse för Vetenskaplig Forskning grant CTS 18-65 and the Kempestiftelserna grant SMK 1860. DAC was supported in part also by funds from the Novo Nordisk Foundation (Grant no. NNF17OC0026486) awarded to Dr. Emmanuelle Charpentier at MIMS, The Laboratory for Molecular Infection Medicine Sweden). AS was funded by MIMS Excellence by Choice Postdoctoral Programme stipend under the patronage of Emmanuelle Charpentier grant SMK-1532.2 and Svenska Sällskapet för Medicinsk Forskning (SSMF) Postdoctoral grant PD20-0022.

The authors are thankful to Yu Wang and Jibin Sun, Tianjin Institute of Industrial Biotechnology, Chinese Academy of Sciences, Tianjin, China for kindly gifting the pnCas9-AID-YU and pgRNA-ccdB plasmids; Javier Pizarro-Cerdá, Institut Pasteur, Paris, France for providing pSU2.1rp-mCherry plasmid; and Justin L. Sonnenburg, Stanford University School of Medicine, Stanford, USA for providing sfGFP carrying *Bacteroides thetaiotaomicron* strain.

Author contributions as per CASRAI are as follows. Conceptualization: AS, AP, DAC.

Formal analysis: AS, ROA, DAC. Funding acquisition: AS, AP, DAC. Investigation: AS, ROA. Methodology: AS, ROA, AP, DAC. Project administration: AP, DAC. Resources: AP, DAC. Supervision: AP, DAC. Visualization: AS, ROA. Writing – original draft: AS. Writing – reviewing and editing: AP, DAC, AS.

## Supplementary Material

**Supplementary Methods S1.** Detailed methodology of construction of plasmids, strains and cloning guide RNA.

**Supplementary Figure S1.** Determination of J23119 promoter activity in *S. flexneri*. (A) The map of pgRNA showing the restriction sites used to clone sfGFP and to obtain pgRNA-(R)-GFP. The sfGFP has a synthetic ribosome binding site and is cloned under the J23119 promoter. (B) *S. flexneri* strains carrying pgRNA-X as non-fluorescent control and pgRNA-(R)-GFP streaked on a TSA plate supplemented with carbenicillin (50 μg/ml).

**Supplementary Figure S2.** *S. flexneri::mCherry* transformants visualized by Typhoon scanner (GE Healthcare) prior to mutagenesis experiment to verify fluorescence.

**Supplementary Figure S3.** (A) Mutation frequency in *E. coli::mCherry*, determined as the ratio of the of number of non-fluorescent (mutant) colonies to the total number of colonies, after 4, 6, and 20 hours of mutagenesis with or without IPTG induction. (B) Titer of *E. coli::mCherry* during mutagenesis determined by counting the colonies spotted after 2, 4, 6, and 20h of mutagenesis with or without IPTG. (C) Mutation frequency in *S. flexneri*::*mCherry*, determined as the ratio of the of number of non-fluorescent (mutant) colonies to the total number of colonies, after 4 and 6 hours of mutagenesis with or without IPTG induction. (D) Titer of *S. flexneri::mCherry* during mutagenesis determined by counting the colonies spotted after 2, 4, and 6 h of mutagenesis with or without IPTG. The numbers on the X-axis in each graph correspond to the conditions represented in (Fig. 2A). Each dot represents a technical replicate and is color-coded to represent a biological replicate. The bars represent the median of three independent experiments. Statistical significance was determined by performing two-way ANOVA followed by Sidák’s multiple comparison tests for of each condition with respect to the control (condition 1) and to determine the significance of for IPTG addition (ns – not significant, \**P*<0.0332, \*\**P*<0.0021, \*\*\**P*<0.0002, \*\*\*\**P*<0.0001).

**Supplementary Figure S4:** (A) Microscopic images of 50:50 mixture of *S. flexneri* WT carrying pgRNA-(R)-GFP and *S. flexneri::mScarlet* carrying control plasmid pgRNA-X cultures taken using different filters. (B) Immunodetection of nCas9-AID fusion protein using anti-Cas9 antibody. The left panel is a representative of multiple Western Blots and the right panel is a representative SDS-PAGE gel of the same samples run in parallel. (C) Mutation frequency in *S. flexneri*, determined as the ratio of the number of nalidixic acid resistant (mutant) colonies to the total number of colonies, after 4 and 6 h of mutagenesis with and without IPTG induction. The numbers on the X-axis correspond to the conditions represented in (Fig. 4A). Each dot represents a technical replicate and is color-coded to represent a biological replicate. In some technical replicates, the mutation frequency was below the technically possible limit of observation (1×10^-9^). The bars represent the median of three independent experiments. Statistical significance was determined by performing two-way ANOVA followed by Sidák’s multiple comparison tests for each condition with respect to the control (condition 1) and for IPTG addition (ns – not significant, \**P*<0.0332, \*\**P*<0.0021, \*\*\**P*<0.0002, \*\*\*\**P*<0.0001). (D) Normalized (to *cysG* and *hcaT*) expression of *icsB, ipgB*, and *ipaC* genes in different mutant strains. Statistical significance was determined by performing (a) a Kruskal-Wallis test (non-parametric) with Dunn’s multiple comparison test and (b) One-way ANOVA (parametric) with Dunnett’s multiple comparison test. The results are represented as (a)/(b) on each bar (ns – not significant, \**P*<0.0332, \*\**P*<0.0021, \*\*\**P*<0.0002, \*\*\*\**P*<0.0001).

**Supplementary Figure S5:** Overview of the workflow to generate multiple parallel mutations in *Shigella* in high throughput. Cloning the spacer oligos in pgRNA and transformation in *E. coli* DH5α is done on Day 1. The transformants obtained can be inoculated to extract plasmids later on Day 2 or overnight cultures can be made for plasmid extraction on Day 3. pgRNA are transformed on Day 3 and *S. flexneri* colonies carrying both plasmids are selected on Day 4 after overnight incubation. Inoculations can be made to carry out mutagenesis later on the same day (Day 4) or overnight cultures can be prepared for mutagenesis on Day 5. Depending on the incubation, colony PCRs can be performed on Day 5 or Day 6 (the day following mutagenesis).

**Supplementary Table S1.** Bacterial strains, cells and plasmids used in this study

**Supplementary Table S2**. Oligonucleotides used in this study

## References

1. Kotloff KL, Nataro JP, Blackwelder WC, Nasrin D, Farag TH, Panchalingam S, Wu Y, Sow SO, Sur D, Breiman RF, Faruque AS, Zaidi AK, Saha D, Alonso PL, Tamboura B, Sanogo D, Onwuchekwa U, Manna B, Ramamurthy T, Kanungo S, Ochieng JB, Omore R, Oundo JO, Hossain A, Das SK, Ahmed S, Qureshi S, Quadri F, Adegbola RA, Antonio M, Hossain MJ, Akinsola A, Mandomando I, Nhampossa T, Acácio S, Biswas K, O’Reilly CE, Mintz ED, Berkeley LY, Muhsen K, Sommerfelt H, Robins-Browne RM, Levine MM. 2013. Burden and aetiology of diarrhoeal disease in infants and young children in developing countries (the Global Enteric Multicenter Study, GEMS): a prospective, case-control study. Lancet 382:209–222.

2. Anderson M, Sansonetti PJ, Marteyn BS. 2016. Shigella Diversity and Changing Landscape: Insights for the Twenty-First Century. Front Cell Infect Microbiol 6:45.

3. Khalil IA, Troeger C, Blacker BF, Rao PC, Brown A, Atherly DE, Brewer TG, Engmann CM, Houpt ER, Kang G, Kotloff KL, Levine MM, Luby SP, MacLennan CA, Pan WK, Pavlinac PB, Platts-Mills JA, Qadri F, Riddle MS, Ryan ET, Shoultz DA, Steele AD, Walson JL, Sanders JW, Mokdad AH, Murray CJL, Hay SI, Reiner RC. 2018. Morbidity and mortality due to shigella and enterotoxigenic Escherichia coli diarrhoea: the Global Burden of Disease Study 1990–2016. The Lancet Infectious Diseases 18:1229–1240.

4. Kotloff KL, Riddle MS, Platts-Mills JA, Pavlinac P, Zaidi AKM. 2018. Shigellosis. The Lancet 391:801–812.

5. Mattock E, Blocker AJ. 2017. How Do the Virulence Factors of Shigella Work Together to Cause Disease? Front Cell Infect Microbiol 7:64.

6. Schnupf P, Sansonetti PJ. 2019. Shigella Pathogenesis: New Insights through Advanced Methodologies. Microbiology Spectrum 7:7.2.28.

7. Cervantes-Rivera R, Tronnet S, Puhar A. 2020. Complete genome sequence and annotation of the laboratory reference strain Shigella flexneri serotype 5a M90T and genome-wide transcriptional start site determination. BMC Genomics 21:285.

8. Datsenko KA, Wanner BL. 2000. One-step inactivation of chromosomal genes in Escherichia coli K-12 using PCR products. PNAS 97:6640–6645.

9. Campbell-Valois F-X, Sachse M, Sansonetti PJ, Parsot C. 2015. Escape of Actively Secreting Shigella flexneri from ATG8/LC3-Positive Vacuoles Formed during Cell-To-Cell Spread Is Facilitated by IcsB and VirA. mBio 6:e02567–02514.

10. Li Z, Liu W, Fu J, Cheng S, Xu Y, Wang Z, Liu X, Shi X, Liu Y, Qi X, Liu X, Ding J, Shao F. 2021. Shigella evades pyroptosis by arginine ADP-riboxanation of caspase-11. Nature 599:290–295.

11. Sidik S, Kottwitz H, Benjamin J, Ryu J, Jarrar A, Garduno R, Rohde JR. 2014. A Shigella flexneri Virulence Plasmid Encoded Factor Controls Production of Outer Membrane Vesicles. G3 (Bethesda) 4:2493–2503.

12. Baba T, Ara T, Hasegawa M, Takai Y, Okumura Y, Baba M, Datsenko KA, Tomita M, Wanner BL, Mori H. 2006. Construction of Escherichia coli K-12 in-frame, single-gene knockout mutants: the Keio collection. Mol Syst Biol 2:2006.0008.

13. Charpentier E, Doudna JA. 2013. Rewriting a genome. Nature 495:50–51.

14. Mali P, Yang L, Esvelt KM, Aach J, Guell M, DiCarlo JE, Norville JE, Church GM. 2013. RNA-guided human genome engineering via Cas9. Science 339:823–826.

15. Bikard D, Euler CW, Jiang W, Nussenzweig PM, Goldberg GW, Duportet X, Fischetti VA, Marraffini LA. 2014. Exploiting CRISPR-Cas nucleases to produce sequence-specific antimicrobials. Nat Biotechnol 32:1146–1150.

16. Jiang W, Bikard D, Cox D, Zhang F, Marraffini LA. 2013. RNA-guided editing of bacterial genomes using CRISPR-Cas systems. 3. Nature Biotechnology 31:233–239.

17. Jiang Y, Chen B, Duan C, Sun B, Yang J, Yang S. 2015. Multigene Editing in the Escherichia coli Genome via the CRISPR-Cas9 System. Appl Environ Microbiol 81:2506–2514.

18. Komor AC, Kim YB, Packer MS, Zuris JA, Liu DR. 2016. Programmable editing of a target base in genomic DNA without double-stranded DNA cleavage. 7603. Nature 533:420–424.

19. Gaudelli NM, Komor AC, Rees HA, Packer MS, Badran AH, Bryson DI, Liu DR. 2017. Programmable base editing of A•T to G•C in genomic DNA without DNA cleavage. Nature 551:464–471.

20. Nishida K, Arazoe T, Yachie N, Banno S, Kakimoto M, Tabata M, Mochizuki M, Miyabe A, Araki M, Hara KY, Shimatani Z, Kondo A. 2016. Targeted nucleotide editing using hybrid prokaryotic and vertebrate adaptive immune systems. Science 353:aaf8729.

21. Banno S, Nishida K, Arazoe T, Mitsunobu H, Kondo A. 2018. Deaminase-mediated multiplex genome editing in Escherichia coli. Nat Microbiol 3:423–429.

22. Wang Y, Liu Y, Liu J, Guo Y, Fan L, Ni X, Zheng X, Wang M, Zheng P, Sun J, Ma Y. 2018. MACBETH: Multiplex automated Corynebacterium glutamicum base editing method. Metabolic Engineering 47:200–210.

23. Bartolomé B, Jubete Y, Martínez E, de la Cruz F. 1991. Construction and properties of a family of pACYC184-derived cloning vectors compatible with pBR322 and its derivatives. Gene 102:75–78.

24. Gay P, Le Coq D, Steinmetz M, Berkelman T, Kado CI. 1985. Positive selection procedure for entrapment of insertion sequence elements in gram-negative bacteria. J Bacteriol 164:918–921.

25. Tobe T, Sasakawa C, Okada N, Honma Y, Yoshikawa M. 1992. vacB, a novel chromosomal gene required for expression of virulence genes on the large plasmid of Shigella flexneri. Journal of Bacteriology 174:6359–6367.

26. Yu H, Wu Z, Chen X, Ji Q, Tao S. 2020. CRISPR-CBEI: a Designing and Analyzing Tool Kit for Cytosine Base Editor-Mediated Gene Inactivation. mSystems 5:e00350–20.

27. Bernard P, Gabant P, Bahassi EM, Couturier M. 1994. Positive-selection vectors using the F plasmid ccdB killer gene. Gene 148:71–74.

28. Dickerson SK, Market E, Besmer E, Papavasiliou FN. 2003. AID Mediates Hypermutation by Deaminating Single Stranded DNA. Journal of Experimental Medicine 197:1291–1296.

29. Kuscu C, Adli M. 2016. CRISPR-Cas9-AID base editor is a powerful gain-of-function screening tool. Nat Methods 13:983–984.

30. Luria SE, Delbrück M. 1943. Mutations of Bacteria from Virus Sensitivity to Virus Resistance. Genetics 28:491–511.

31. Ghosh A, N. S, Saha S. 2020. Survey of drug resistance associated gene mutations in Mycobacterium tuberculosis, ESKAPE and other bacterial species. Sci Rep 10:8957.

32. Allaoui A, Sansonetti PJ, Parsot C. 1993. MxiD, an outer membrane protein necessary for the secretion of the Shigella flexneri lpa invasins. Mol Microbiol 7:59–68.

33. Sharma A, Puhar A. 2019. Gentamicin Protection Assay to Determine the Number of Intracellular Bacteria during Infection of Human TC7 Intestinal Epithelial Cells by Shigella flexneri. Bio-protocol 9:e3292–e3292.

34. Bahrani FK, Sansonetti PJ, Parsot C. 1997. Secretion of Ipa proteins by Shigella flexneri: inducer molecules and kinetics of activation. Infect Immun 65:4005–4010.

35. Bernardini ML, Mounier J, d’Hauteville H, Coquis-Rondon M, Sansonetti PJ. 1989. Identification of icsA, a plasmid locus of Shigella flexneri that governs bacterial intra-and intercellular spread through interaction with F-actin. Proc Natl Acad Sci U S A 86:3867–3871.

36. Carpenter CD, Cooley BJ, Needham BD, Fisher CR, Trent MS, Gordon V, Payne SM. 2014. The Vps/VacJ ABC Transporter Is Required for Intercellular Spread of Shigella flexneri. Infection and Immunity 82:660–669.

37. Oaks EV, Wingfield ME, Formal SB. 1985. Plaque formation by virulent Shigella flexneri. Infect Immun 48:124–129.

38. Sharma A, Puhar A. 2019. Plaque Assay to Determine Invasion and Intercellular Dissemination of Shigella flexneri in TC7 Human Intestinal Epithelial Cells. Bio-protocol 9:e3293–e3293.

39. Allaoui A, Mounier J, Prévost M-C, Sansonetti PJ, Parsot C. 1992. icsB: a Shigella flexneri virulence gene necessary for the lysis of protrusions during intercellular spread. Molecular Microbiology 6:1605–1616.

40. Rathman M, Jouirhi N, Allaoui A, Sansonetti P, Parsot C, Tran Van Nhieu G. 2000. The development of a FACS-based strategy for the isolation of Shigella flexneri mutants that are deficient in intercellular spread. Molecular Microbiology 35:974–990.

41. Hachani A, Biskri L, Rossi G, Marty A, Ménard R, Sansonetti P, Parsot C, Van Nhieu GT, Bernardini ML, Allaoui A. 2008. IpgB1 and IpgB2, two homologous effectors secreted via the Mxi-Spa type III secretion apparatus, cooperate to mediate polarized cell invasion and inflammatory potential of Shigella flexenri. Microbes and Infection 10:260–268.

42. Graffeuil A, Guerrero-Castro J, Assefa A, Uhlin BE, Cisneros DA. 2022. Polar mutagenesis of polycistronic bacterial transcriptional units using Cas12a. Microb Cell Fact 21:139.

43. Fels U, Gevaert K, Van Damme P. 2020. Bacterial Genetic Engineering by Means of Recombineering for Reverse Genetics. Frontiers in Microbiology 11:2161.

44. Brochado AR, Typas A. 2013. High-throughput approaches to understanding gene function and mapping network architecture in bacteria. Current Opinion in Microbiology 16:199–206.

45. Freed NE, Bumann D, Silander OK. 2016. Combining Shigella Tn-seq data with gold-standard E. coli gene deletion data suggests rare transitions between essential and non-essential gene functionality. BMC Microbiology 16:203.

46. Wang L, Xue W, Yan L, Li X, Wei J, Chen M, Wu J, Yang B, Yang L, Chen J. 2017. Enhanced base editing by co-expression of free uracil DNA glycosylase inhibitor. Cell Res 27:1289–1292.

47. Bridges BA. 2001. Hypermutation in bacteria and other cellular systems. Phil Trans R Soc Lond B 356:29–39.

48. Sternberg SH, Redding S, Jinek M, Greene EC, Doudna JA. 2014. DNA interrogation by the CRISPR RNA-guided endonuclease Cas9. Nature 507:62–67.

49. Mateus A, Shah M, Hevler J, Kurzawa N, Bobonis J, Typas A, Savitski MM. 2021. Transcriptional and Post-Transcriptional Polar Effects in Bacterial Gene Deletion Libraries. mSystems 6:e00813–21.

50. Hove-Jensen B. 2008. Two-step method for curing Escherichia coli of ColE1-derived plasmids. Journal of Microbiological Methods 72:208–213.

51. Wang T, Wei JJ, Sabatini DM, Lander ES. 2014. Genetic screens in human cells using the CRISPR-Cas9 system. Science 343:80–84.

52. Kleinstiver BP, Prew MS, Tsai SQ, Topkar VV, Nguyen NT, Zheng Z, Gonzales APW, Li Z, Peterson RT, Yeh J-RJ, Aryee MJ, Joung JK. 2015. Engineered CRISPR-Cas9 nucleases with altered PAM specificities. Nature 523:481–485.

53. Povolotskaya IS, Kondrashov FA, Ledda A, Vlasov PK. 2012. Stop codons in bacteria are not selectively equivalent. Biology Direct 7:30.

54. McKenzie GJ, Craig NL. 2006. Fast, easy and efficient: site-specific insertion of transgenes into Enterobacterial chromosomes using Tn7 without need for selection of the insertion event. BMC Microbiol 6:39.

55. Tadala L, Langenbach D, Dannborg M, Cervantes-Rivera R, Sharma A, Vieth K, Rieckmann LM, Wanders A, Cisneros DA, Puhar A. 2022. Infection-induced membrane ruffling initiates danger and immune signaling via the mechanosensor PIEZO1. Cell Reports 40:111173.

56. García-Nafría J, Watson JF, Greger IH. 2016. IVA cloning: A single-tube universal cloning system exploiting bacterial In Vivo Assembly. Sci Rep 6:27459.

57. Inoue H, Nojima H, Okayama H. 1990. High efficiency transformation of Escherichia coli with plasmids. Gene 96:23–28.

58. Clerc P, Sansonetti PJ. 1987. Entry of Shigella flexneri into HeLa cells: evidence for directed phagocytosis involving actin polymerization and myosin accumulation. Infect Immun 55:2681–2688.

59. 2008. Cytoplasmic targeting of IpaC to the bacterial pole directs polar type III secretion in Shigella. The EMBO Journal 27:447–457.

60. Cervantes-Rivera R, Puhar A. 2020. Whole-genome Identification of Transcriptional Start Sites by Differential RNA-seq in Bacteria. Bio-protocol 10:e3757–e3757.

61. Zhou K, Zhou L, Lim Q’En, Zou R, Stephanopoulos G, Too H-P. 2011. Novel reference genes for quantifying transcriptional responses of Escherichia coli to protein overexpression by quantitative PCR. BMC Molecular Biology 12:18.

