## Supplementary figures for "CRISPR-Cas guided mutagenesis of chromosome and virulence plasmid in *Shigella flexneri* by cytosine base editing"

### Figure S1

**A**

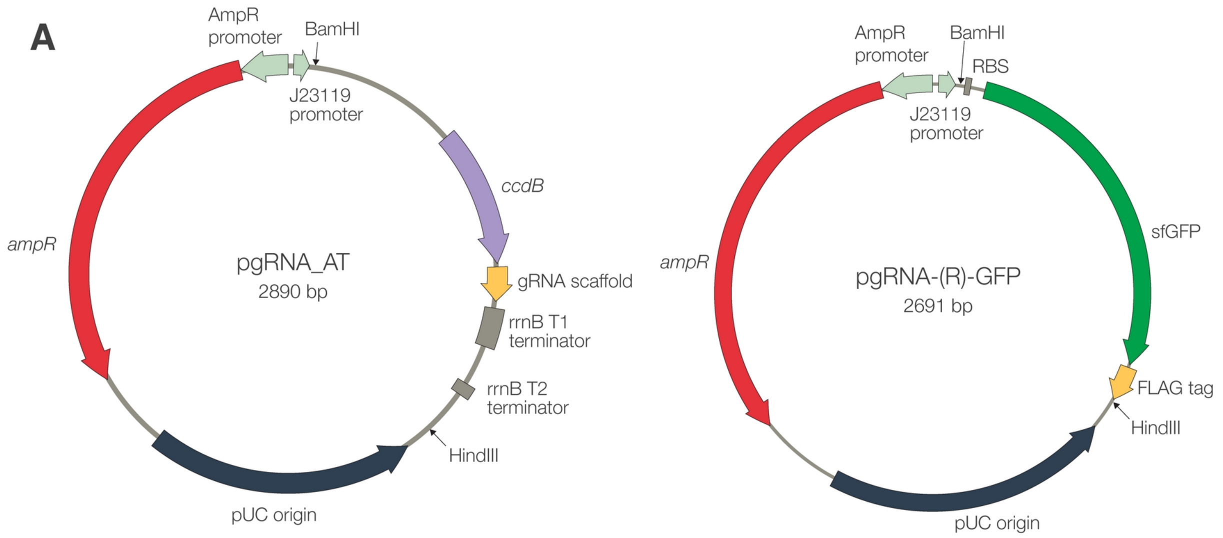
AmpR promoter
BamHI
J23119 promoter
ccdB
ampR
pgRNA_AT
2890 bp
gRNA scaffold
rrnB T1 terminator
rrnB T2 terminator
HindIII
pUC origin
AmpR promoter
BamHI
RBS
J23119 promoter
sfGFP
ampR
pgRNA-(R)-GFP
2691 bp
FLAG tag
HindIII
pUC origin

**B**

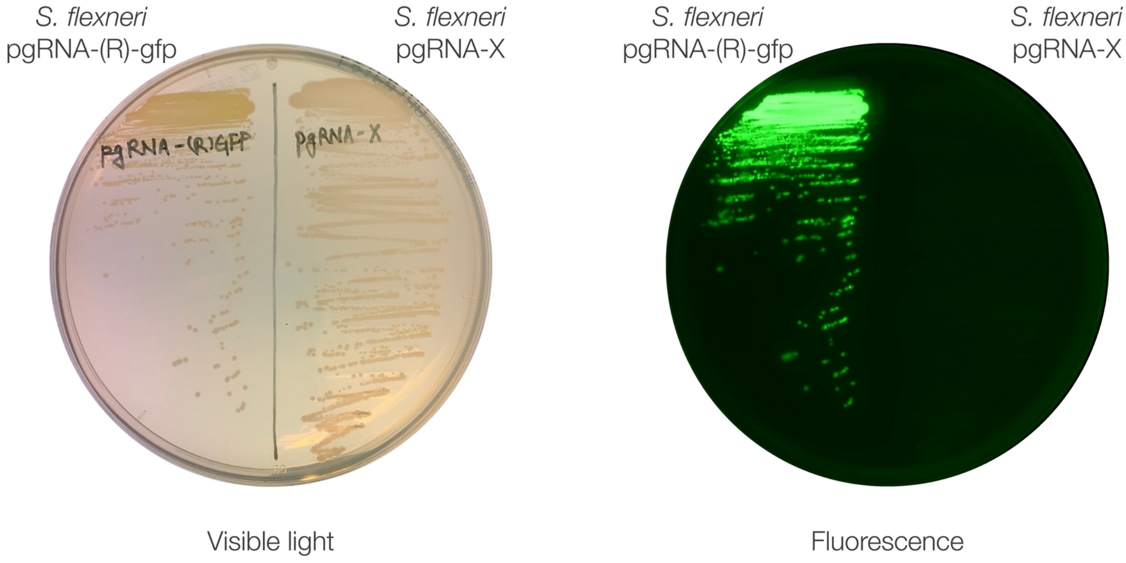
S. flexneri pgRNA-(R)-gfp
S. flexneri pgRNA-X
S. flexneri pgRNA-(R)-gfp
S. flexneri pgRNA-X
Visible light
Fluorescence

### Figure S2

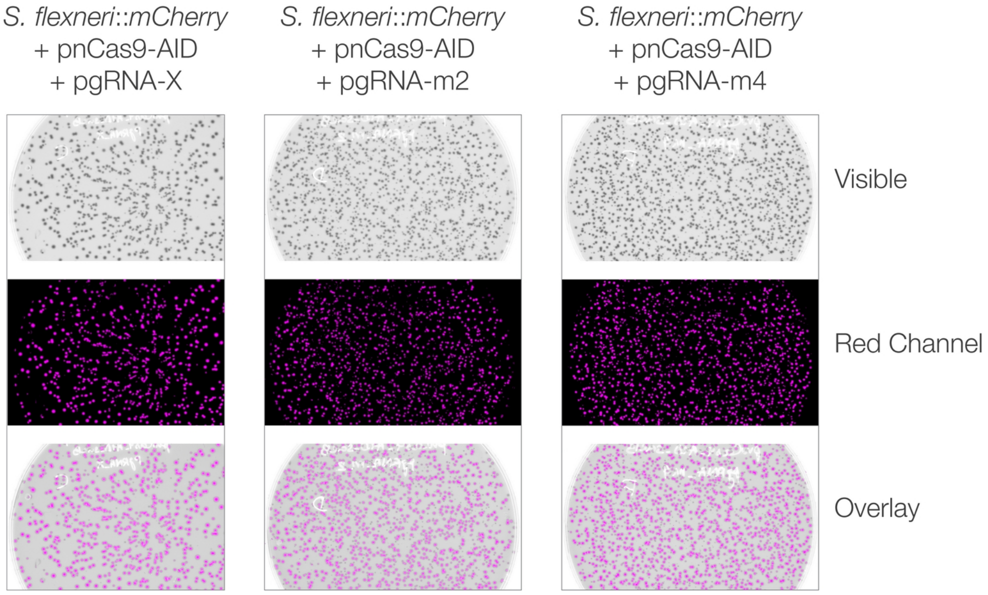
*S. flexneri*::*mCherry*
+ pnCas9-AID
+ pgRNA-X
*S. flexneri*::*mCherry*
+ pnCas9-AID
+ pgRNA-m2
*S. flexneri*::*mCherry*
+ pnCas9-AID
+ pgRNA-m4
Visible
Red Channel
Overlay

### Figure S3

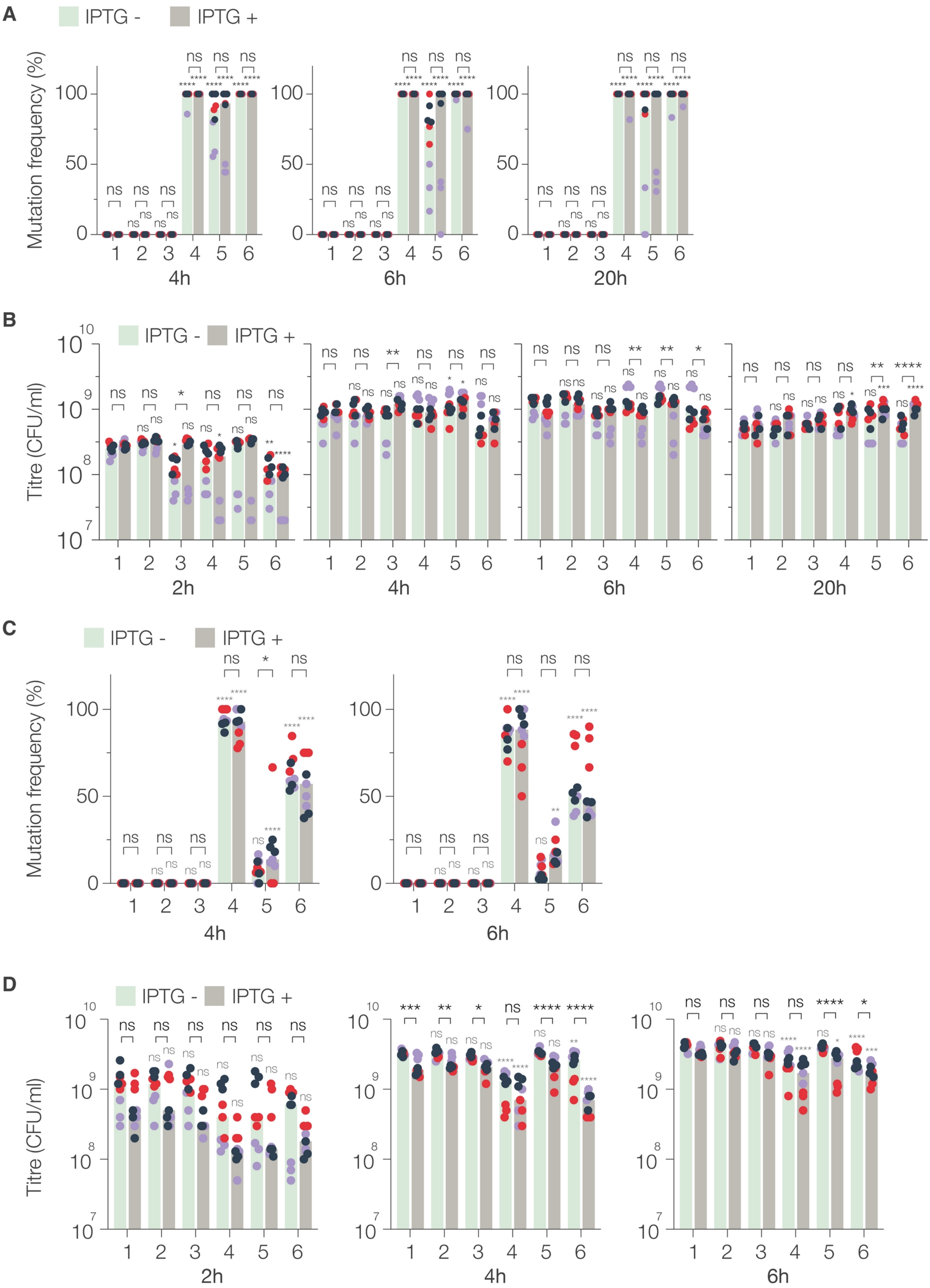
A
IPTG -
IPTG +
Mutation frequency (%)
4h
6h
20h
B
Titre (CFU/ml)
2h
4h
6h
20h
C
Mutation frequency (%)
4h
6h
D
Titre (CFU/ml)
2h
4h
6h

### Figure S4

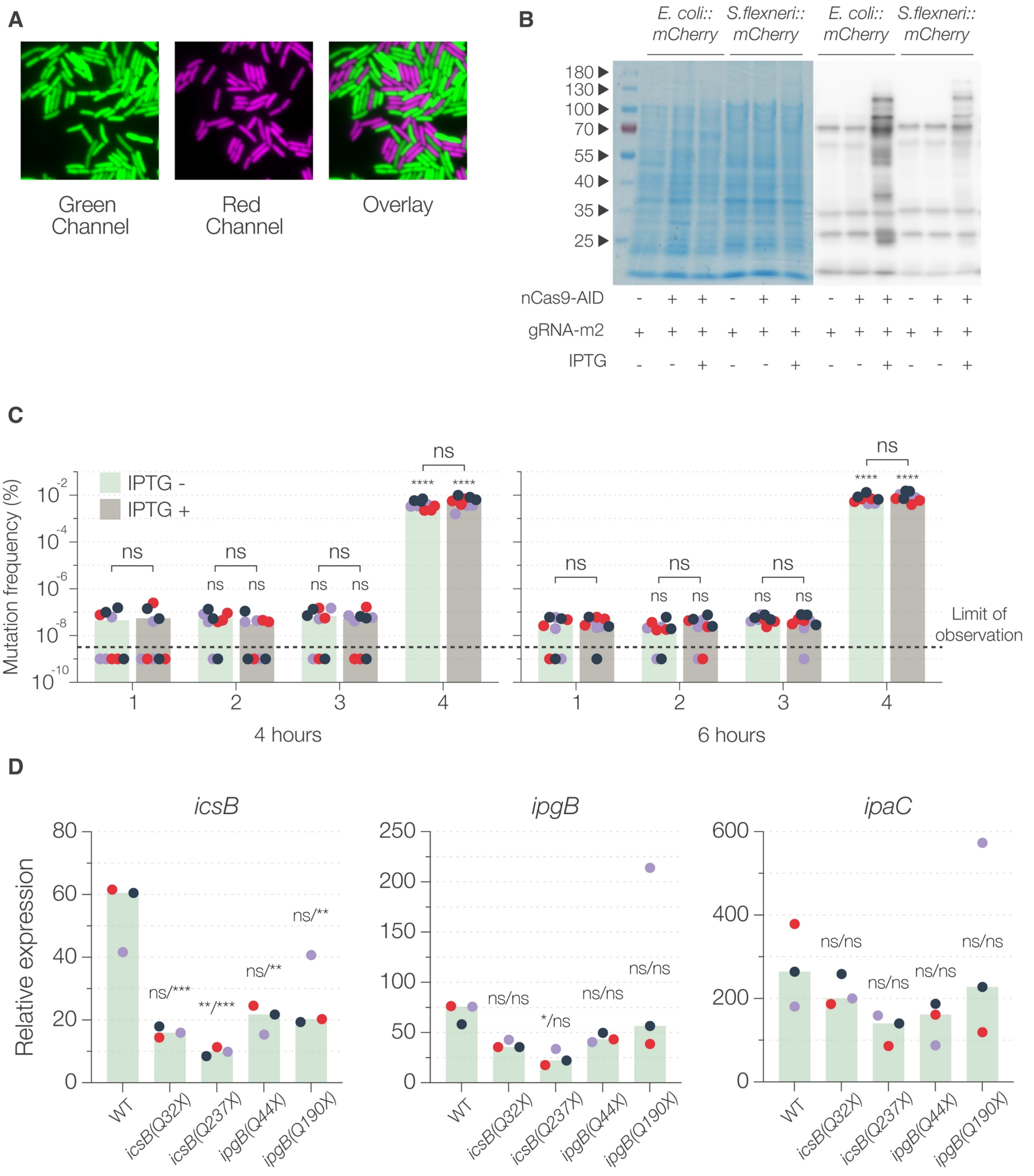
A
Green Channel
Red Channel
Overlay
B
E. coli:: mCherry
S.flexneri:: mCherry
E. coli:: mCherry
S.flexneri:: mCherry
180
130
100
70
55
40
35
25
nCas9-AID
gRNA-m2
IPTG
C
IPTG -
IPTG +
Mutation frequency (%)
ns
****
Limit of observation
4 hours
6 hours
D
icsB
ipgB
ipaC
Relative expression
WT
icsB(Q32X)
icsB(Q237X)
ipgB(Q44X)
ipgB(Q190X)
ns/***
**/***
ns/**
ns/ns
*/ns

### Figure S5

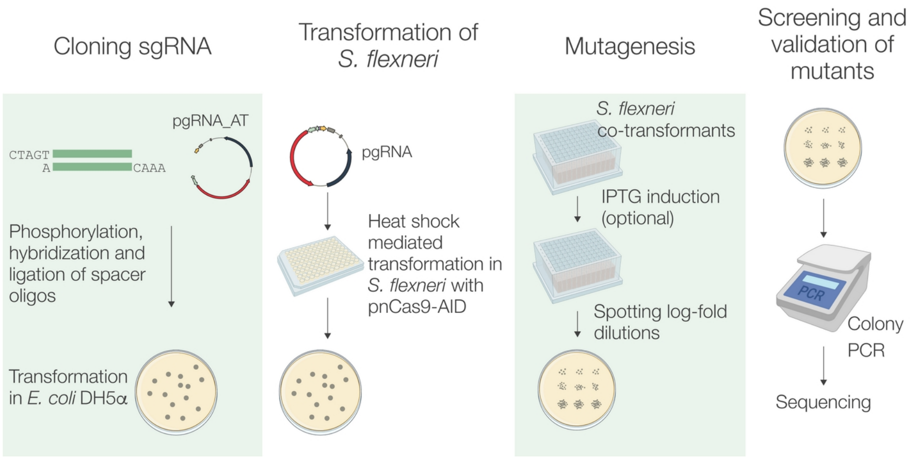

Cloning sgRNA
Transformation of
*S. flexneri*
Mutagenesis
Screening and
validation of
mutants
pgRNA_AT
CTAGT
A
CAAA
Phosphorylation,
hybridization and
ligation of spacer
oligos
Transformation
in *E. coli* DH5α
pgRNA
Heat shock
mediated
transformation in
*S. flexneri* with
pnCas9-AID
*S. flexneri*
co-transformants
IPTG induction
(optional)
Spotting log-fold
dilutions
PCR
Colony
PCR
Sequencing

### Supplementary Methods S1

#### Construction of fluorescent strains

**F**luorescent strains of *E. coli* and *S. flexneri*, constitutively expressing *mCherry* and *mScarlet-i* (only *S. flexneri*) from the chromosome, were constructed using Tn7-mediated transposition (1, 2). The plasmid pGRG25 expresses Tn7 transposase which stably inserts a single copy of genes cloned between the repeats of the transposon. The insertion event, downstream the *glmS* gene, has no impact on *S. flexneri* virulence (assessed by gentamicin protection assay, data not shown). *mScarlet-i* was amplified from pQE-60NA-mScarlet-2I using the oligos AT_0157 and AT_0158. The PCR product was digested with BamHI and HindIII and ligated to similarly digested pSU2.1rp-cerulean plasmid, yielding pSU-mScarlet plasmid. *mCherry* and *mScarlet-i* expression cassette (the ORF along with a constitutive promoter and ribosome binding site) was amplified from the plasmid pSU2.1rp-mCherry (3), and pSU-mScarlet, respectively, using the primers AT_0017 and AT_0018. The purified PCR product was digested with NotI and XhoI restriction enzymes and ligated to similarly digested pGRG25 plasmid. The resulting plasmids, pGRG-mCherry or pGRG-mScarlet, were transformed into *E. coli* MG1655 and *S. flexneri* M90T 5a, respectively. The transformants were selected on LA (for *E. coli*) or TSA (for *S. flexneri*) plates supplemented with carbenicillin (100 μg/ml) at 30°C. A single transformant colony was grown overnight in LB or TSB supplemented with carbenicillin at 30°C (since pGRG plasmids have a temperature sensitive pSC101 origin). The overnight cultures were diluted in fresh medium containing carbenicillin and the bacteria were grown till $OD_{600}$ = 0.2 at 30°C. Transposition was induced by the addition of 0.5% L-arabinose and the cultures were grown overnight at 30°C. Dilutions of these cultures were spread on agar plates with no antibiotics and incubated overnight at 37°C, since pGRG plasmids cannot replicate at this temperature. The resulting colonies were checked for fluorescence and the fluorescent colonies were patched onto agar plates containing carbenicillin to ensure the loss of pGRG-mCherry (or pGRG-mScarlet). *S. flexneri* colonies were also patched onto Congo red plates (TSA supplemented with 0.01% Congo red) to select only for the virulent colonies. The chromosomal integration of *mCherry* and *mScarlet-i* was verified by sequencing the PCR product obtained by using primers AT_0059 and AT_0061 for *E. coli*, and primers AT_0062 and AT_0063 for *S. flexneri*.

**Plasmid construction**

The sgRNA expression plasmid (pgRNA_AT) was generated by combining parts from pgRNA-ccdB and a pBluescript SK variant that lacks BsaI site in the β-lactamase gene (4). Using the primers AT_0001 and AT_0002, *ccdB* gene and the downstream guide RNA scaffold were amplified from pgRNA_ccdB. The AT_0001 has a synthetic constitutive promoter, J23119, at the 5’ end, making the resulting PCR product a complete sgRNA expression cassette. Using the primers AT_0004 and AT_0005, the plasmid backbone (origin and resistance marker) was amplified from pBluescript SK. Both these PCR products were digested with EcoRI and HindIII and ligated. The resulting chimeric plasmid was amplified using the primers AT_0087 and AT_088 and the PCR product was phosphorylated and self-ligated to form the pgRNA_AT plasmid. The sgRNA cassette in pgRNA_AT, as a result, has an extra C residue downstream the BsaI restriction site to ensure accurate cloning of spacer oligos. *E. coli* DB3.1 was used as a cloning host for generating pgRNA_AT as it is resistant to the action of *ccdB*. To verify the activity of J23119 promoter in *Shigella*, sfGFP was cloned under the promoter in pgRNA_AT. sfGFP encoding gene was amplified from the genomic DNA of sfGFP expressing *Bacteroides thetaiotaomicron* (5) using the primers AT_0057 and AT_0058. The PCR product was digested with BamHI and HindIII and ligated to similarly digested pgRNA_AT to yield pgRNA-(R)-GFP

(Supplementary Fig. S1). The plasmid was transformed into WT *S. flexneri* and the transformants were selected on TSA plates supplemented with carbenicillin. A single transformant colony of *S. flexneri* carrying pgRNA-X and pgRNA-(R)-GFP was streaked on a TSA plate containing carbenicillin. The plates were incubated overnight at 37°C and imaged using ChemiDoc (Bio-Rad).

**T**he nCas9-AID expressing plasmid, pnCas9-AID, was generated by combining parts of pSU19 and pnCas9-AID-YU by *in-vivo* assembly (IVA) cloning (4, 6). From pnCas9-AID_YU, the nCas9-AID encoding gene, with *tac* promoter and the LacI encoding gene were amplified using the primers AT_0043 and AT_0044. The p15A origin and antibiotic resistance marker were amplified from pSU19 using the primers AT_0008 and AT_0009. Both PCR products were treated with DpnI (to digest the parent plasmids) and an equimolar mixture of both PCR products was transformed into ultracompetent *E. coli* DH5α. The resulting plasmid, named pAS_004, was sequenced to verify the construction. A *sacB* expression cassette was amplified from pYC1000-eforRED plasmid and cloned in the NotI site of pAS_004. The resulting plasmid was designated as pnCas9-AID.

#### Guide RNA design and cloning

**T**he guide RNA for mutagenesis were designed using the program CRISPR_CBEI which detects the mutable sites and provides the sequences of spacer oligos (7). The editing window was set from -16 to -20 from an 'NGG' PAM. We gave preference to mutable sites in the first 50% of the target gene sequence to maximize the odds of creating loss-of-function mutations. The location of the target sequence within the gene can be easily determined from the output of the CRISPR_CBEI program. Multiple mutable sites were found for *mCherry*, *icsA*, *icsB*, and *vacJ* in the first half of the sequence. In case of *ipgB*, the second mutable site was very close to the Cterminus. We still included in the experiments to see whether the phenotype of both mutants was similar. In case of *mxiD*, however, only one mutable site was found, which was located in the second half of the sequence (but resulted in loss-of-function mutants). For every spacer, two complementary oligos were ordered with the forward oligo carrying 5' – CTAGT and the reverse oligo carrying 5' – AAAC and a 3' – A residues. The forward and reverse oligos were annealed in an annealing buffer (10 mM Tris-HCl pH 7.5, 50 mM NaCl, and 1 mM EDTA) by incubating at 95°C for 3 min and letting the mixture cool down to room temperature gradually. The double stranded spacer oligos were 5' phosphorylated using T4 DNA T4 Polynucleotide Kinase (Thermo Fisher Scientific) and ligated to BsaI digested pgRNA_AT using T4 DNA Ligase (Thermo Fisher Scientific). The ligation product was transformed into *E. coli* DH5α, to obtain target-specific sgRNA plasmids. These plasmids were verified by DNA sequencing and named according to the sgRNA they express (e.g. pgRNA_m2 expresses the sgRNA_m2).

**Statistical analysis**

All statistical analyses were performed in GraphPad Prism. The type of analysis and test of significance is mentioned in the respective figure legends. The default P value system of GraphPad Prism was used in all cases and the key is described in the figure legends.

#### References (only for Supplementary Methods S1)

1. McKenzie GJ, Craig NL. 2006. Fast, easy and efficient: site-specific insertion of transgenes into Enterobacterial chromosomes using Tn7 without need for selection of the insertion event. BMC Microbiol 6:39.

2. Tadala L, Langenbach D, Dannborg M, Cervantes-Rivera R, Sharma A, Vieth K, Rieckmann LM, Wanders A, Cisneros DA, Puhar A. 2022. Infection-induced membrane ruffling

initiates danger and immune signaling via the mechanosensor PIEZO1. Cell Reports 40:111173.

3. Campbell-Valois F-X, Sachse M, Sansonetti PJ, Parsot C. 2015. Escape of Actively Secreting Shigella flexneri from ATG8/LC3-Positive Vacuoles Formed during Cell-To-Cell Spread Is Facilitated by IcsB and VirA. mBio 6:e02567-02514.

4. Wang Y, Liu Y, Liu J, Guo Y, Fan L, Ni X, Zheng X, Wang M, Zheng P, Sun J, Ma Y. 2018. MACBETH: Multiplex automated Corynebacterium glutamicum base editing method. Metabolic Engineering 47:200–210.

5. Whitaker WR, Shepherd ES, Sonnenburg JL. 2017. Tunable Expression Tools Enable Single-Cell Strain Distinction in the Gut Microbiome. Cell 169:538-546.e12.

6. García-Nafría J, Watson JF, Greger IH. 2016. IVA cloning: A single-tube universal cloning system exploiting bacterial In Vivo Assembly. Sci Rep 6:27459.

7. Yu H, Wu Z, Chen X, Ji Q, Tao S. 2020. CRISPR-CBEI: a Designing and Analyzing Tool Kit for Cytosine Base Editor-Mediated Gene Inactivation. mSystems 5:e00350-20.

**Supplementary Table S1.** Bacterial strains, cells and plasmids used in this study.

| **Name** | **Description** | **Reference** |
|---|---|---|
| **Bacteria** | | |
| *E. coli* DH5α | General cloning host | |
| *E. coli* DB3.1 | *ccdB* resistant strain, cloning host for generating pgRNA_AT | |
| *E. coli* MG1655 | WT *E. coli* strain | |
| *E. coli*::*mCherry* | *E. coli* MG1655 *attTn7::mCherry*; *E. coli* MG1655 expressing *mCherry* from the chromosome after integration of *mCherry* at the attTn7 site | This study |
| *S. flexneri* M90T 5a | WT *Shigella flexneri* strain | (1) |
| *S. flexneri*::*mCherry* | *S. flexneri attTn7::mCherry*; *S. flexneri* expressing *mCherry* from the chromosome after integration of *mCherry* at the *attTn7* site | (2) |
| *S. flexneri::mScarlet* | *S. flexneri attTn7::mScarlet*; *S. flexneri* expressing *mScarlet-i* from the chromosome after integration of *mScarlet-i* at the *attTn7* site | This study |
| *S. flexneri mxiD(Q323X)* | *S. flexneri mxiD* loss-of-function mutant generated by substituting Gln$_{323}$ by STOP codon (C$_{964}$→T) | This study |
| *S. flexneri icsA(Q59X)* | *S. flexneri icsA* loss-of-function mutant generated by substituting Gln$_{59}$ by STOP codon (C$_{175}$→T) | This study |
| *S. flexneri icsB(Q32X)* | *S. flexneri icsB* loss-of-function mutant generated by | This study |

| | | |
|---|---|---|
| | substituting $Gln_{32}$ by STOP codon ($C_{94}$→T) | |
| *S. flexneri icsB(Q237X)* | *S. flexneri icsB* loss-of-function mutant generated by substituting $Gln_{237}$ by STOP codon ($C_{709}$→T) | This study |
| *S. flexneri ipgB(Q44X)* | *S. flexneri ipgB* loss-of-function mutant generated by substituting $Gln_{44}$ by STOP codon ($C_{130}$→T) | This study |
| *S. flexneri ipgB(Q190X)* | *S. flexneri ipgB* loss-of-function mutant generated by substituting $Gln_{190}$ by STOP codon ($C_{568}$→T) | This study |
| *S. flexneri vacJ(Q90X)* | *S. flexneri vacJ* loss-of-function mutant generated by substituting $Gln_{90}$ by STOP codon ($C_{268}$→T) | This study |
| **Cells** | | |
| TC7 cells | Intestinal epithelial cells | (3) |
| **Plasmids** | | |
| pgRNA_ccdB | sgRNA expression plasmid used in *C. glutamicum* | (4) |
| pgRNA_AT | sgRNA expression plasmid for cloning spacers | This study |
| pBluescript SK | General cloning vector used as a source of backbone for pgRNA_AT | |
| pnCas9-AID-YU | nCas9-AID expressing plasmid used in *C. glutamicum* | (4) |
| pnCas9-AID | nCas9-AID expressing plasmid; carries *sacB* that aids in curing | This study |
| pSU19 | General cloning vector used as a source of backbone for pnCas9-AID | (5) |
| pAS_004 | Intermediate nCas9-AID plasmid before cloning of | This study |

| | | |
|---|---|---|
| | *sacB* expression cassette | |
| pYC1000-eforRED | Source of *sacB* expression cassette | (6) |
| pQE-60NA-mScarlet-2I | Source of *mScarlet-i* | (7) |
| pSU2.1rp-mCherry | Source of *mCherry* expression cassette | (8) |
| pSU2.1rp-cerulean | Plasmid used for cloning *mScarlet-i* (to have the promoter and RBS similar to *mCherry* expression cassette) | (8) |
| pGRG25 | Plasmid for Tn7 transposition | (9) |
| pGRG-mCherry | pGRG25 plasmid with *mCherry* expression cassette cloned for transposition | (2) |
| pGRG-mScarlet | pGRG25 plasmid with *mScarlet-i* expression cassette cloned for transposition | This study |
| pgRNA-(R)-GFP | sgRNA expression plasmid modified to express sfGFP with an RBS | This study |
| pgRNA-X | sgRNA expression plasmid carrying non-targeting spacer | This study |
| pgRNA-m2 | sgRNA expression plasmid carrying mCh2 spacer | This study |
| pgRNA-m3 | sgRNA expression plasmid carrying mCh3 spacer | This study |
| pgRNA-m4 | sgRNA expression plasmid carrying mCh4 spacer | This study |
| pgRNA-icsA_G1 | sgRNA expression plasmid carrying icsA_G1 spacer | This study |
| pgRNA-icsA_G2 | sgRNA expression plasmid carrying icsA_G2 spacer | This study |
| pgRNA_icsA_G3 | sgRNA expression plasmid carrying icsA_G3 spacer | This study |
| pgRNA-icsB_G1 | sgRNA expression plasmid carrying icsB_G1 spacer | This study |

| | | |
|---|---|---|
| pgRNA-icsB_G2 | sgRNA expression plasmid carrying icsB_G2 spacer | This study |
| pgRNA-mxiD_G1 | sgRNA expression plasmid carrying mxiD_G1 spacer | This study |
| pgRNA-vacJ_G1 | sgRNA expression plasmid carrying vacJ_G1 spacer | This study |
| pgRNA-vacJ_G2 | sgRNA expression plasmid carrying vacJ_G2 spacer | This study |
| pgRNA-vacJ_G3 | sgRNA expression plasmid carrying vacJ_G3 spacer | This study |

References

1. Sansonetti PJ, Kopecko DJ, Formal SB. 1982. Involvement of a plasmid in the invasive ability of Shigella flexneri. Infect Immun 35:852–860.

2. Tadala L, Langenbach D, Dannborg M, Cervantes-Rivera R, Sharma A, Vieth K, Rieckmann LM, Wanders A, Cisneros DA, Puhar A. 2022. Infection-induced membrane ruffling initiates danger and immune signaling via the mechanosensor PIEZO1. Cell Reports 40:111173.

3. Chantret I, Rodolosse A, Barbat A, Dussaulx E, Brot-Laroche E, Zweibaum A, Rousset M. 1994. Differential expression of sucrase-isomaltase in clones isolated from early and late passages of the cell line Caco-2: evidence for glucose-dependent negative regulation. J Cell Sci 107 ( Pt 1):213–225.

4. Wang Y, Liu Y, Liu J, Guo Y, Fan L, Ni X, Zheng X, Wang M, Zheng P, Sun J, Ma Y. 2018. MACBETH: Multiplex automated Corynebacterium glutamicum base editing method. Metabolic Engineering 47:200–210.

5. Bartolomé B, Jubete Y, Martínez E, de la Cruz F. 1991. Construction and properties of a family of pACYC184-derived cloning vectors compatible with pBR322 and its derivatives. Gene 102:75–78.

6. Yan M-Y, Yan H-Q, Ren G-X, Zhao J-P, Guo X-P, Sun Y-C. 2017. CRISPR-Cas12a-Assisted Recombineering in Bacteria. Appl Environ Microbiol 83:e00947-17.

7. Valbuena FM, Fitzgerald I, Strack RL, Andruska N, Smith L, Glick BS. 2020. A photostable monomeric superfolder green fluorescent protein. Traffic 21:534–544.

8. Campbell-Valois F-X, Sachse M, Sansonetti PJ, Parsot C. 2015. Escape of Actively Secreting Shigella flexneri from ATG8/LC3-Positive Vacuoles Formed during Cell-To-Cell Spread Is Facilitated by IcsB and VirA. mBio 6:e02567-02514.

9. McKenzie GJ, Craig NL. 2006. Fast, easy and efficient: site-specific insertion of transgenes into Enterobacterial chromosomes using Tn7 without need for selection of the insertion event. BMC Microbiol 6:39.

**Supplementary Table S2**. Oligonucleotides used in this study

| Name | Sequence | Purpose |
|---|---|---|
| AT_0001 | ATTAGAATTCTTGACAGCTAGCTCAGTCCTAGGTATAATACTAGTGAGACCACGCGTGGATCC | Amplification of gRNA expression cassette from pgRNA_ccdB (ref), the forward primer introduces J23119 promoter |
| AT_0002 | GATCAAGCTTGGAGGTCGAAGCCGCACG | |
| AT_0004 | GTACAAGCTTTGAGCAAAAGGCCAGCAAAAG | Amplification of the plasmid backbone from pBlueScript II |
| AT_0005 | ATCGGAATTCAATGTGCGCGGAACCCC | |
| AT_0008 | GCATCCTAGGTGCTTTTGCCGTTACGCAC | Amplification of the plasmid backbone from pSU19 |
| AT_0009 | ATTAGCGGCCGCTAATTGCGTTGCGCTCACTG | |
| AT_0017 | TTATGCGGCCGCGCGTTGGCCGATTC | Amplification of *mCherry* cassette from pSU2.1rp-mCherry |
| AT_0018 | ATTACTCGAGCCAGGGTTTTCCCAGTC | |
| AT_0043 | GCGCAACGCAATTAGCGGCCGCTAATGCATCCGGGGCTGATCCCCG | Amplification of dCas9-AID from pCas9-AID (ref) for IVA cloning |
| AT_0044 | GCGTAACGGCAAAAGCACCTAGGATGCGGCCATCCGTCAGGATGGCC | |
| AT_0057 | ATTAGGATCCGAAAGAGGAGAAATACTAGATGGATTCAAAAAAATTTAAAATAATGCG | Cloning *gfp* (with RBS) in pgRNA |
| AT_0058 | TAGCAAGCTTAACCTTTATCATCATCGTCC | |
| AT_0059 | TTCCTGGGCCGTGGCGATC | Sequencing and verification of *E. coli*::*mCherry* |
| AT_0061 | TACCCTGGTAGTTAACTTTATTACCGG | |
| AT_0062 | CCGTTCCGCTGCAGCTG | Sequencing and verification of |

| | | |
|---|---|---|
| | | *S. flexneri*::*mCherry* |
| AT_0063 | ACACAGCCACTGGATCGG | |
| AT_0087 | CCGTTTTAGAGCTAGAAATAGCAAG | Modification of the pgRNA cloning site by adding a C residue |
| AT_0088 | AGACCTTATATTCCCCAGAACATCAG | |
| AT_0093 | CTAGTTCAGTTCATGTACGGCTCCA | Cloning mCh2 spacer in pgRNA |
| AT_0094 | AAACGTGGAGCCGTACATGAACTGA | |
| AT_0095 | CTAGTGACCCAGGACTCCTCCCTGC | Cloning mCh3 spacer in pgRNA |
| AT_0096 | AAACGCAGGGAGGAGTCCTGGGTCA | |
| AT_0097 | CTAGTGCTCCCACTTGAAGCCCTCG | Cloning mCh4 spacer in pgRNA |
| AT_0098 | AAACGCGAGGGCTTCAAGTGGGAGC | |
| AT_0099 | CTAGTGGATCCACTAGTGCGGCCGC | Cloning gRNA-X in pgRNA |
| AT_0100 | AAACGGCGGCCGCACTAGTGGATCC | |
| AT_0105 | CTAGTCATAAACCGCCGAGTCACCA | Cloning gyrA spacers in pgRNA |
| AT_0106 | AAACTGGTGACTCGGCGGTTTATGA | |
| AT_0109 | CTAGTTGGCAAGGAACGGCTTCATT | Cloning mxiD_G1 spacers in pgRNA |
| AT_0110 | AAACAATGAAGCCGTTCCTTGCCAA | |
| AT_0111 | CTAGTGACCCAATGTTCACAGGGGG | Cloning icsA_G1 spacers in pgRNA |
| AT_0112 | AAACCCCCCTGTGAACATTGGGTCA | |
| AT_0113 | CTAGTCAATGTTCACAGGGGGGGGC | Cloning icsA_G2 spacers in pgRNA |
| AT_0114 | AAACGCCCCCCCCTGTGAACATTGA | |
| AT_0115 | CTAGTTCAAGAACTTCATTTTTCAG | Cloning icsA_G3 spacers in |
| AT_0116 | AAACCTGAAAAATGAAGTTCTTGAA | |

| | | |
|---|---|---|
| | | pgRNA |
| AT_0119 | CTAGTCTTGCAGGGCGACCCTTATC | Cloning vacJ_G1 spacers in pgRNA |
| AT_0120 | AAACGATAAGGGTCGCCCTGCAAGA | |
| AT_0121 | CTAGTTTGCAGGGCGACCCTTATCA | Cloning vacJ_G2 spacers in pgRNA |
| AT_0122 | AAACTGATAAGGGTCGCCCTGCAAA | |
| AT_0123 | CTAGTTGCAGGGCGACCCTTATCAG | Cloning vacJ_G3 spacers in pgRNA |
| AT_0124 | AAACCTGATAAGGGTCGCCCTGCAA | |
| AT_0137 | CTAGTAGCACAGAAATTCAACCTAA | Cloning icsB_G1 spacers in pgRNA |
| AT_0138 | AAACTTAGGTTGAATTTCTGTGCTA | |
| AT_0141 | CTAGTAATCAAAAAAAAGACCCCTA | Cloning icsB_G2 spacers in pgRNA |
| AT_0142 | AAACTAGGGGTCTTTTTTTTGATTA | |
| AT_0143 | ACATATGCTCACGAGGTACA | Sequencing *icsB* |
| AT_0144 | CCATACCAGCACAGTTTTC | |
| AT_0147 | ATTGCATATCCAGAAACC | Sequencing *mxiD* |
| AT_0148 | TAGCCGGAATATTCTCTTG | |
| AT_0151 | GGAACTACGCTTCTGGTG | Sequencing *vacJ* |
| AT_0152 | CGTGAAGCTACCGTAGAAC | |
| AT_0153 | AGGTAAATTTCTCCCGTTG | Sequencing *icsA* |
| AT_0154 | TGATTACAGAGAGGCTGCT | |
| gRNA-X | GGATCCACTAGTGCGGCCGC | Random non-binding spacer |
| mCh2 | TCAGTTCATGTACGGCTCCA | *mCherry* spacer for $C_{205}$→T, $Gln_{69}$→STOP mutation |
| mCh3 | GACCCAGGACTCCTCCCTGC | *mCherry* spacer for $C_{340}$→T, |

| | | |
|---|---|---|
| | | $Gln_{114}$→STOP mutation |
| mCh4 | GCTCCCACTTGAAGCCCTCG | *mCherry* spacer for $G_{294}$→A, $Trp_{98}$→STOP mutation |
| icsA_G1 | GACCCAATGTTCACAGGGGG | *icsA* spacer for $C_{37}$→T, $Gln_{13}$→STOP mutation |
| icsA_G2 | CAATGTTCACAGGGGGGGGC | *icsA* spacer for $C_{175}$→T, $Gln_{59}$→STOP mutation |
| icsA_G3 | TCAAGAACTTCATTTTTCAG | *icsA* spacer for $C_{442}$→T, $Gln_{148}$→STOP mutation |
| icsB_G1 | AGCACAGAAATTCAACCTAA | *icsB* spacer for $C_{94}$→T, $Gln_{32}$→STOP mutation |
| icsB_G2 | AATCAAAAAAAAGACCCCTA | *icsB* spacer for $C_{709}$→T, $Gln_{237}$→STOP mutation |
| mxiD_G1 | TGGCAAGGAACGGCTTCATT | *mxiD* spacer for $C_{964}$→T, $Gln_{323}$→STOP mutation |
| vacJ_G1 | CTTGCAGGGCGACCCTTATC | *vacJ* spacer for $C_{268}$→T, $Gln_{90}$→STOP mutation |
| vacJ_G2 | TTGCAGGGCGACCCTTATCA | *vacJ* spacer for $C_{268}$→T, $Gln_{90}$→STOP mutation |
| vacJ_G3 | TGCAGGGCGACCCTTATCAG | *vacJ* spacer for $C_{268}$→T, $Gln_{90}$→STOP mutation |
| sgRNA- | CATAAACCGCCGAGTCACCA | *gyrA* spacer for $G_{259}$→A, $Asp_{87}$→Asn mutation |

| |
|---|
| gyrA |

2

### Tables

**Table 1. Details of characterized *S. flexneri* genes that were selected for mutagenesis.**

The selected genes are found both on the virulence plasmid and the chromosome. The variation in the number of spacers and their corresponding mutable sites is also mentioned.

| Gene | Location | Spacers | Function | Mutant phenotype | Reference |
|---|---|---|---|---|---|
| *vacJ* | Chromosome | Three spacers; same mutable site | Lysis of protrusion during intracellular dissemination | Dissemination defect | (36) |
| *icsA* | Virulence plasmid | Two spacers; different mutable sites | Actin-based motility | Dissemination defect | (35) |
| *icsB* | Virulence plasmid | Two spacers; different mutable sites | Post-invasion virulence factor | Dissemination defect (?) | (39, 40) |
| *mxiD* | Virulence plasmid | One spacer | Structural component of the injectisome | Invasion defect | (32) |
